# Reversal of ageing- and injury-induced vision loss by Tet-dependent epigenetic reprogramming

**DOI:** 10.1101/710210

**Authors:** Yuancheng Lu, Anitha Krishnan, Benedikt Brommer, Xiao Tian, Margarita Meer, Daniel L. Vera, Chen Wang, Qiurui Zeng, Doudou Yu, Michael S. Bonkowski, Jae-Hyun Yang, Emma M. Hoffmann, Songlin Zhou, Ekaterina Korobkina, Noah Davidsohn, Michael B. Schultz, Karolina Chwalek, Luis A. Rajman, George M. Church, Konrad Hochedlinger, Vadim N. Gladyshev, Steve Horvath, Meredith S. Gregory-Ksander, Bruce R. Ksander, Zhigang He, David A. Sinclair

## Abstract

Ageing is a degenerative process leading to tissue dysfunction and death. A proposed cause of ageing is the accumulation of epigenetic noise, which disrupts youthful gene expression patterns that are required for cells to function optimally and recover from damage^1–3^. Changes to DNA methylation patterns over time form the basis of an ‘ageing clock’^4, 5^, but whether old individuals retain information to reset the clock and, if so, whether this would improve tissue function is not known. Of all the tissues in the body, the central nervous system (CNS) is one of the first to lose regenerative capacity^6, 7^. Using the eye as a model tissue, we show that expression of Oct4, Sox2, and Klf4 genes (OSK) in mice resets youthful gene expression patterns and the DNA methylation age of retinal ganglion cells, promotes axon regeneration after optic nerve crush injury, and restores vision in a mouse model of glaucoma and in normal old mice. This process, which we call <u>re</u>co<u>v</u>ery of information <u>v</u>ia <u>e</u>pigenetic <u>r</u>eprogramming or REVIVER, requires the DNA demethylases Tet1 and Tet2, indicating that DNA methylation patterns don’t just indicate age, they participate in ageing. Thus, old tissues retain a faithful record of youthful epigenetic information that can be accessed for functional age reversal.

---

The metaphor of the epigenetic landscape, first invoked by Waddington to explain embryonic development^8, 9^, is increasingly seen as relevant to the other end of life^9^. Evidence from yeast and mammals supports an Information Theory of Ageing, in which the loss of epigenetic information disrupts youthful gene expression patterns^1–3^, leading to cellular dysfunction and senescence.^10^

In mammals, progressive DNA methylation changes serve as an epigenetic clock, but whether they are merely an effect or a driver of ageing is not known^4, 5^. In cell culture, the ectopic expression of the four Yamanaka transcription factors, namely Oct4, Sox2, Klf4, and c-Myc (OSKM)^11^, can reprogram somatic cells to become pluripotent stem cells, a process that erases most DNA methylation marks and leads to the loss of cellular identity^4, 12^. *In vivo*, ectopic, transgene-mediated expression of these four genes alleviates progeroid symptoms in a mouse model of Hutchison-Guilford Syndrome, indicating that OSKM might counteract normal ageing^13^. Continual expression of all four factors, however, induces teratomas^14^ or causes death within days^13^, ostensibly due to tissue dysplasia^15^.

Ageing is generally considered a unidirectional process akin an increase in entropy, but living systems are open, not closed, and in some cases can fully reset biological age, examples being “immortal” cnidarians and the cloning of animals by nuclear transfer^16^. Having previously found evidence for epigenetic noise as an underlying cause of ageing^2, 3^, we wondered whether mammalian cells might retain a faithful copy of epigenetic information from earlier in life, analogous to Shannon’s “observer” system in Information Theory, essentially a back-up copy of the original signal to allow for its reconstitution at the receiving end if information is lost or noise is introduced during transmission^17^.

To test this hypothesis, we introduced the expression of three-gene OSK combination in fibroblasts from old mice and measured its effect on RNA levels of genes known to be altered with age, including H2A, H2B, LaminB1, and Chaf1b. We excluded the c-Myc gene from these experiments because it is an oncogene that reduces lifespan^18^. OSK-treated old fibroblasts promoted youthful gene expression patterns, with no apparent loss of cellular identity or the induction of Nanog, an early embryonic transcription factor that can induce teratomas (Extended Data Fig.1a-c).

**Figure 1.**
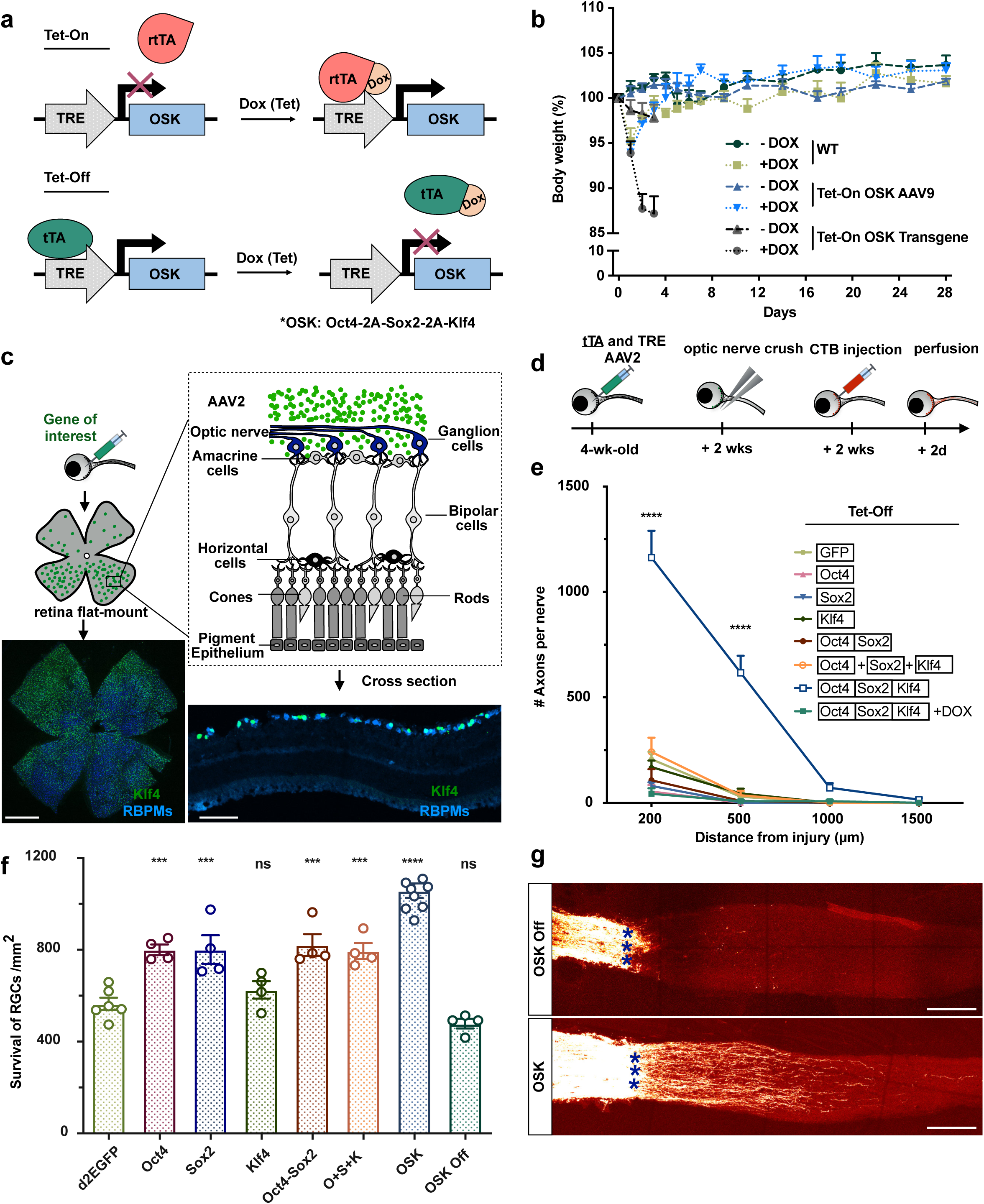
AAV-delivered polycistronic OSK is non-toxic and induces CNS axon regeneration. **a**, Schematic of the Tet-On and Tet-Off AAV vectors used in this study. **b**, Body weight of WT mice, OSK transgenic mice and AAV9-mediated OSK-expressing mice (1.0 x 10^12^ gene copies) with or without doxycycline induction in the first 4 weeks (n = 5, 3, 6, 4, 6 and 3, respectively). **c**, Schematic of intravitreal AAV2 injection to target retina ganglion cells. A representative retinal whole mount and retinal cross section 2 weeks after intravitreal injection of OSK AAV. Stains for Klf4 (green) and RBPMS (blue) show the transduction efficiency and targeted nerve fiber layer. Scales bars = 1 mm and 100 µm respectively. **d**, Experimental outline of the optic nerve crush study using the Tet-Off system. Alexa-conjugated (555 nm) cholera toxin subunit B (CTB) was used for anterograde axonal tracing. **e**, Quantification of regenerating axons 16 days after crush injury at multiple distances distal to the lesion site in mice treated with d2EGFP, Oct4, Sox2, Klf4, Oct4-Sox2, O+S+K on separate vectors, or OSK AAV2. Error bars indicate s.e.m. (n = 5, 4, 4, 4, 4, 4, 7, 5). **f**, Survival of RBPMS-positive cells in the RGC layer transduced with different AAV2 16 days after crush injury (n = 4-8). **g**, Representative images of longitudinal sections through the optic nerve showing CTB-labeled regenerative axons on day 16 post-injury in wild-type mice with an intravitreal injection of AAV2-tTA and AAV2-TRE-OSK in the presence or absence of Dox. Blue asterisks indicate the crush site. Scale bars = 200 µm. ***, P<0.001, ****, P<0.0001, one-way ANOVA with Bonferroni correction in **e**, **f**, relative to d2EGFP in **f**.

Next, we tested if a similar restoration was possible in mice. To deliver and control OSK expression *in vivo*, we engineered a tightly regulated adeno-associated viral (AAV) vector under the control of tetracycline response element (TRE) promoter (Fig.1a)^19, 20^. The TRE promoter can be activated either by rtTA in the presence of DOX (the “Tet-ON” system) or by tTA in the absence of DOX (“Tet-OFF”). Extraneous AAV sequences were removed so the vector could accommodate OSK as a poly-cistron. To test if ectopic OSK expression was toxic *in vivo*, we infected 5 month-old C57BL/6J mice with AAV9-rtTA and AAV9-TRE-OSK delivered intravenously (1.0 x 10^12^ gene copies total), then induced OSK expression by providing doxycycline in the drinking water (Extended Data Fig. 2a). Surprisingly, continuous induction of OSK for over a year had no discernable negative effect on the mice (Fig.1b and Extended Data Fig. 2b)^20^, ostensibly because we avoided high-level expression in the intestine (Extended Data Fig.2c-e), thus avoiding the dysplasia and weight loss seen in other studies^15^.

**Figure 2.**
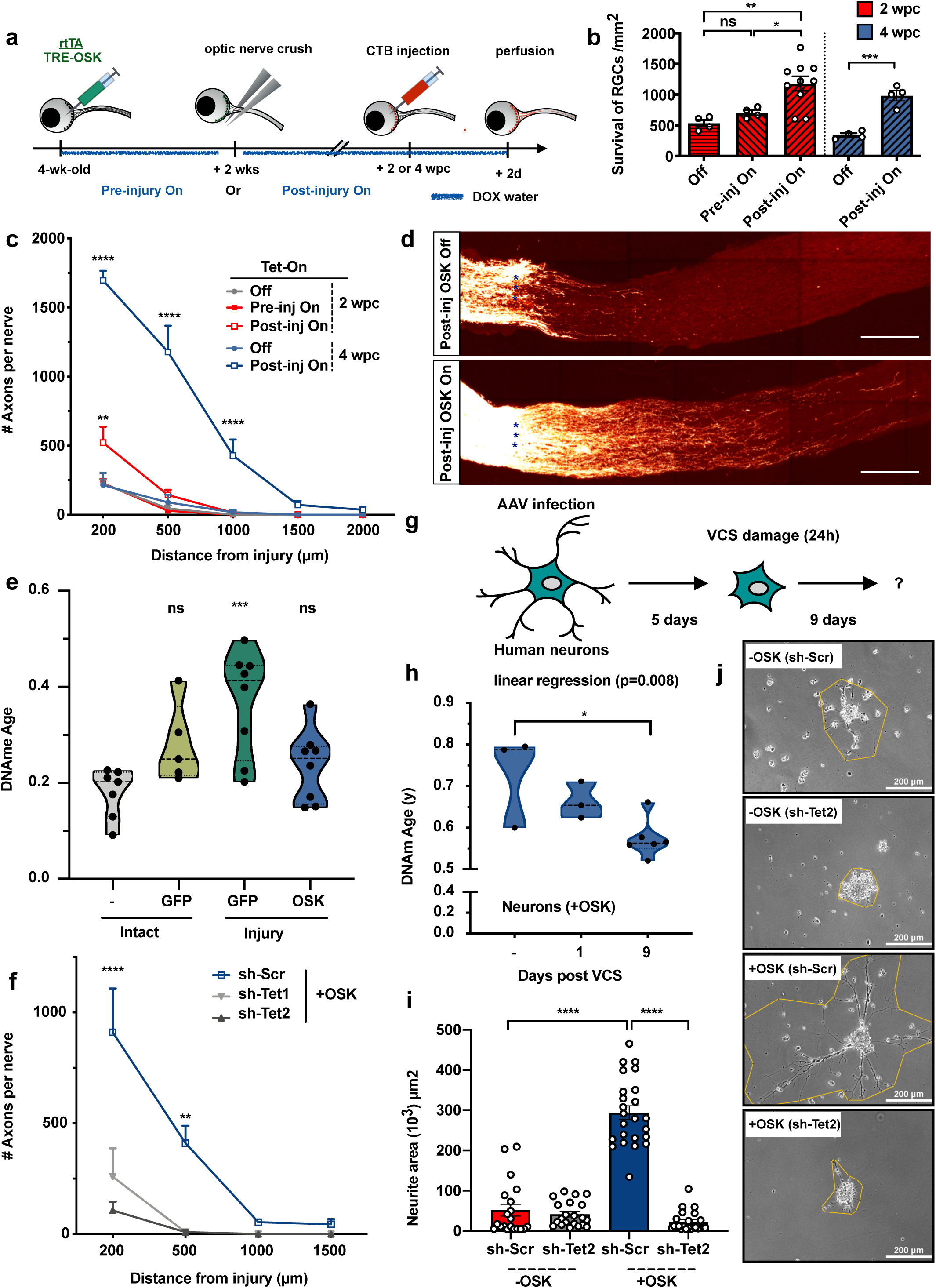
OSK expression promotes axon regeneration and RGC survival through a Tet-dependent mechanism. **a**, Experimental strategies for pre- and post-injury induction of OSK expression (wpc - weeks post crush). **b,** RGC survival in response to OSK induction pre- or post-injury. **c,** Axon regeneration in response to OSK induction pre- or post-injury. **d,** Representative longitudinal sections through the optic nerve showing CTB-labeled axons 4 weeks after crush injury, with or without post-injury induction of OSK. Blue asterisks indicate optic nerve crush site. Scale bars represent 200 µm. **e,** rDNA methylation age of 1-month-old RGCs isolated from axon-intact retinas infected with or without GFP, or from axon-injured retinas infected with GFP-AAV or OSK-AAV 4 days after nerve crush. **f,** Axons regeneration in retinas co-transduced with AAV2 vectors encoding polycistronic OSK, tTA, and shRNA vectors with a scrambled sequence (Scr), Tet1, or Tet2 sequences to knockdown Tet DNA dioxygenases/demethylases. **g,** Axon regeneration in human neurons post-vincristine (VCS) damage. **h,** DNA methylation age of human neurons with OSK expression pre-damage (Day -) or after VCS damage (Day 1 and 9), estimated by skin and blood cell clocks. **i,** Neurite area in each AAV treatment group. **j,** Representative images of human neurons and the neurite area 9 days post-VCS damage. *p < 0.05, **p < 0.01, **** p < 0.0001, one-way ANOVA with Bonferroni’s multiple comparison test in **b**, **c**, **e**, **f**, **h**, **i**, relative to first group in **e.** Linear regression p value in **h** refers to a decrease in DNAme Age.

Almost all species experience a decline in regenerative potential during ageing. In mammals, one of the first systems to lose its regenerative potential is the CNS.^6^ Retinal ganglion cells (RGCs) are part of the CNS that project axons away from the retina towards the brain, forming the optic nerve. During embryogenesis and in neonates, RGCs can regenerate if damaged, but this capacity is lost within days after birth^7^. Then, throughout adulthood, the function of these cells continues to decline^21^. To date, attempts to reverse acute or chronic damage to the CNS have been moderately successful, and no treatments have successfully restored eyesight.

To test whether it is possible to provide adult RGCs with the regenerative capacity, we induced OSK in an optic nerve crush injury model. The Tet-Off system carrying OSK, either in separate AAVs or in the same AAV, was injected into the vitreous body, resulting in efficient, selective, and doxycycline-responsive gene expression in RGCs (Fig.1c). As a negative control, a group of mice were also continuously treated with doxycycline to repress OSK expression (Extended Data Fig. 3a). Two weeks after AAV delivery, we performed optic nerve crush. Axon length and optic nerve density were determined 16 days later (Fig.1d).

**Figure 3.**
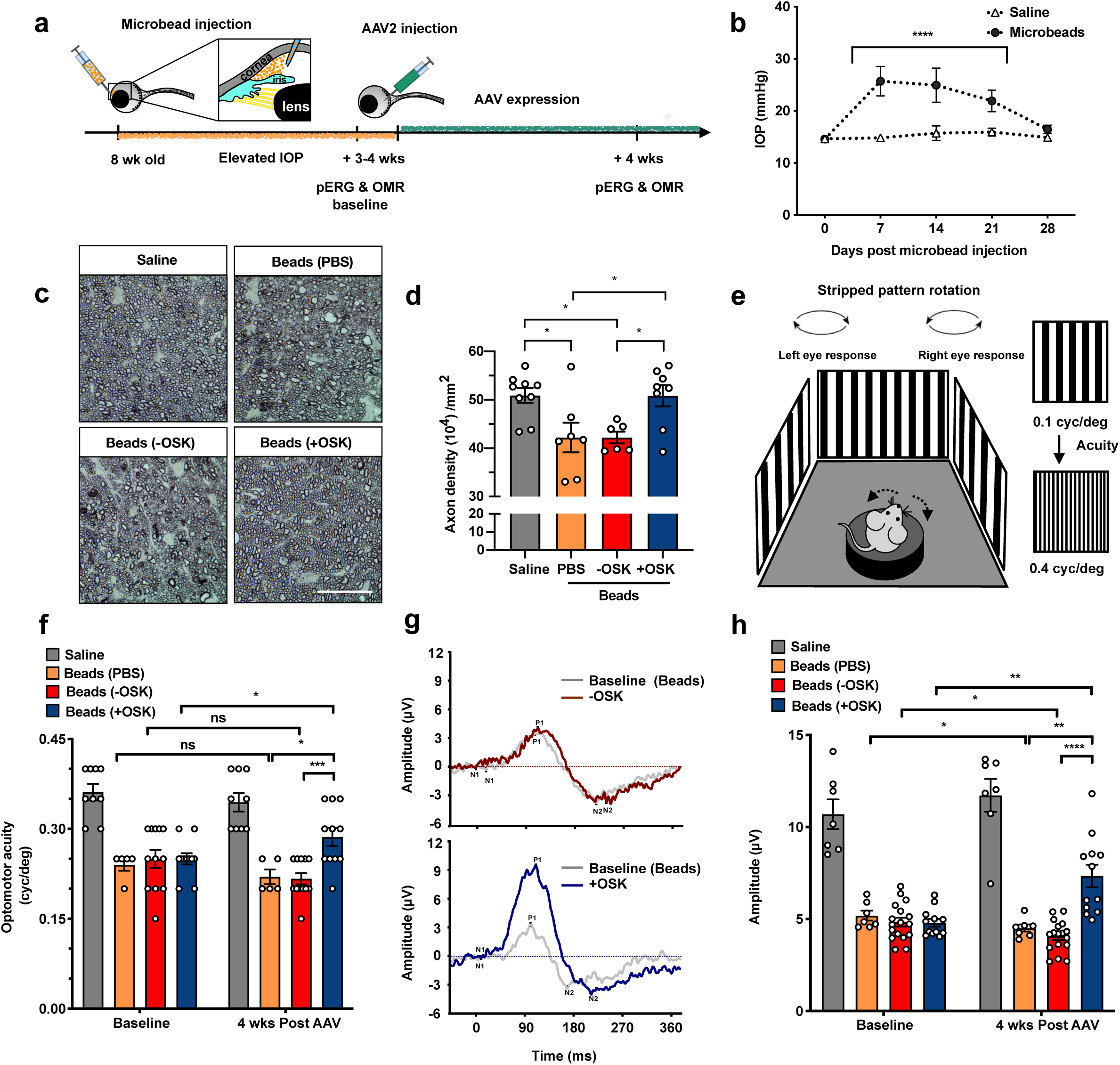
OSK AAV treatment restores visual function in an experimental model of glaucoma. **a**, Experimental outline. **b,** Intraocular pressure (IOP) measured weekly by rebound tonometry for the first 4 weeks post-microbead injection. **c**, Representative micrographs of PPD-stained optic nerve cross-sections at 4 weeks after AAV2 or PBS injection. Scale bars = 50 µm. -OSK: AAV-rtTA+AAV-TRE-OSK; +OSK: AAV-tTA+AAV-TRE-OSK. **d**, Quantification of healthy axons in optic nerves 4 wks after PBS or AAV injection. **e**, High-contrast visual stimulation to measure optomotor response. To assess vision, reflexive head movement was tracked in response to the rotation of a moving stripe pattern that increases in spatial frequency. **f,** Spatial frequency threshold response of each mouse measured before treatment and 4 weeks after intravitreal injection of AAV vectors. **g**, Representative PERG waveforms recorded from the same eye at baseline before AAV injection and four weeks later after -OSK (upper graph) or +OSK AAV (lower graph) treatment. **h**, Mean PERG amplitudes of recordings measured from each mouse at baseline before treatment and 4 weeks after intravitreal injection of AAVs. \**P* < 0.05; \*\**P* < 0.01; \*\*\**P* < 0.001, \*\*\*\**P* < 0.0001. Two-way ANOVA with Bonferroni correction between groups was used for the overall effect of time and treatment. A paired t-test was used to compare before and after treatments.

The greatest extent of axon regeneration and RGC survival, independent of RGC proliferation (Extended Data Fig.4a), occurred when all three genes were delivered via the same AAV as a polycistron (Fig. 1e-g). Indeed, when polycistronic OSK was induced for 12-16 weeks, regenerating RGC axon fibers extended over 5 mm into the chiasm, where optic nerve connects to brain (Extended Data Fig.4b, c). When genes were co-delivered by separate AAVs, no effect on axon regeneration was observed, ostensibly due to the lower frequency of co-infection (Extended Data Fig.3c, d). When delivered singly, OCT4 and SOX2 alone increased RGC survival slightly, but none of the single factors alone had any effect on regenerative capacity (Fig. 1e, f). Because Klf4 has been reported to repress rather than promote axonal growth^22, 23^, we also tested a dual-cistron of just Oct4 and Sox2, but observed no regenerative effect even in the absence of Klf4 (Fig. 1e, f).

Utilizing the Tet-On AAV system for its rapid on-rate (Extended Data Fig. 3b and 5a, b), we tested the effect of inducing OSK expression before or after damage. Significant improvement in axon regeneration only occurred when OSK expression was induced after injury; the longer the duration of OSK induction post-injury, the greater the distance the axons extended, with no increase in the total number of RGCs (Fig. 2b-d). By comparing infected RGCs numbers pre- and post-injury, survival rate of OSK infected RGC was ∼2.5-3.0 times that of uninfected or GFP-infected RGCs (52 vs. 17%-20%) (Extended Data Fig. 5c, d), indicating OSK’s protective and regenerative effects are largely cell-intrinsic. The PTEN-mTOR-S6K pathway, previously shown to improve RGC survival and axon regeneration *in vivo*^24^, was not activated in OSK-infected cells post-injury (Extended Data Fig. 6a, b), indicating a new pathway might be involved.

Given the necessity of post-injury OSK expression and the known role of Yamanaka factors to reverse DNA methylation (DNAme) age during partial or fully reprogramming *in vitro*^4, 12, 25^, we wondered whether neuronal injury advanced epigenomic age and whether OSK’s benefits were due to the preservation of a younger epigenome. Genomic DNA from FACS-isolated RGCs was obtained from retinas that are intact or 4-days after axonal injury in the presence or absence of OSK induction and subjected reduced-representation bisulfite sequencing (RRBS). A newly published rDNAme clock^26^ provided the best site coverage (70/72 CpG sites) relative to other available mouse clocks^27, 28^, and its age estimate remained highly correlated with chronological age of RGCs (Extended Data Fig. 7a and Methods). Consistent with the hypothesis, in the absence of global methylation changes, injured RGCs experienced an acceleration of the epigenetic clock and OSK expression counteracted this effect (Fig. 2e and Extended Data Fig. 7b).

Ten-Eleven-Translocation (TET) dioxygenases are known for their ability to remove DNA demethylation at CpG sites. Because Yamanaka factors promote *in vitro* reprogramming by upregulating Tet1 and Tet2, but have no effect on Tet3^29, 30^, we tested whether Tet1 and Tet2 were required for the beneficial effects of OSK on RGCs. We utilized previously well characterized AAVs expressing short-hairpin RNAs against Tet1 and Tet2 (sh-Tet1 and sh-Tet2)^31–33^ and validated their high co-transduction rate (near 70%) with OSK AAV in the RGCs (Extended Data Fig. 6c, d). Knockdown of either Tet1 or Tet2 blocked the ability of OSK to promote RGC survival and regeneration (Fig. 2f and Extended Data Fig. 6e).

To explore if neuronal reprogramming might be applicable to humans, we performed axon regeneration assays on differentiated SH-SY5Y human neuronal cultures (Fig. 2g), with and without OSK induction (Extended Data Fig. 8a, b). Similar to results of mouse RGCs *in vivo* (Extended Data Fig. 4a), OSK did not induce human neuron cell proliferation (Extended Data Fig. 8c, d). Axon degeneration was then induced by a 24 hr treatment with vincristine (VCS), a chemotherapeutic agent, and cells were then allowed to recover for 9 days. Again, we measured the DNA methylation age of AAV-transduced neurons using a clock for *in vitro* studies^34^. Paralleling the RGCs, DNA methylation age was significantly increased after VCS-induced damage of human neurons (Extended Data Fig. 8e), and OSK expression not only prevented this increase but also restored a younger DNA methylation age without a global reduction of DNA methylation (Fig.2h and Extended Data Fig. 7c). At Day 9 post-damage, the neurite area was 15-fold greater in the rejuvenated OSK-transduced cells than controls (Extended Data Fig. 8f, g). The recovery from damage was completely blocked by validated Tet2 knockdown (Fig. 2i, j, Extended Data Fig. 8h) even in presence of high OSK expression (Extended Data Fig. 8i), but was not dependent on mTOR-S6K pathway (Extended Data Fig. 8j, k), again paralleling mouse RGCs. Thus, the ability of OSK to reprogram neurons and promote axon growth is a conserved, DNA demethylation-dependent cell intrinsic process. We refer to this process as the <u>re</u>co<u>v</u>ery of information <u>v</u>ia <u>e</u>pigenetic <u>r</u>eprogramming, or “REVIVER” for short.

**Figure 4.**
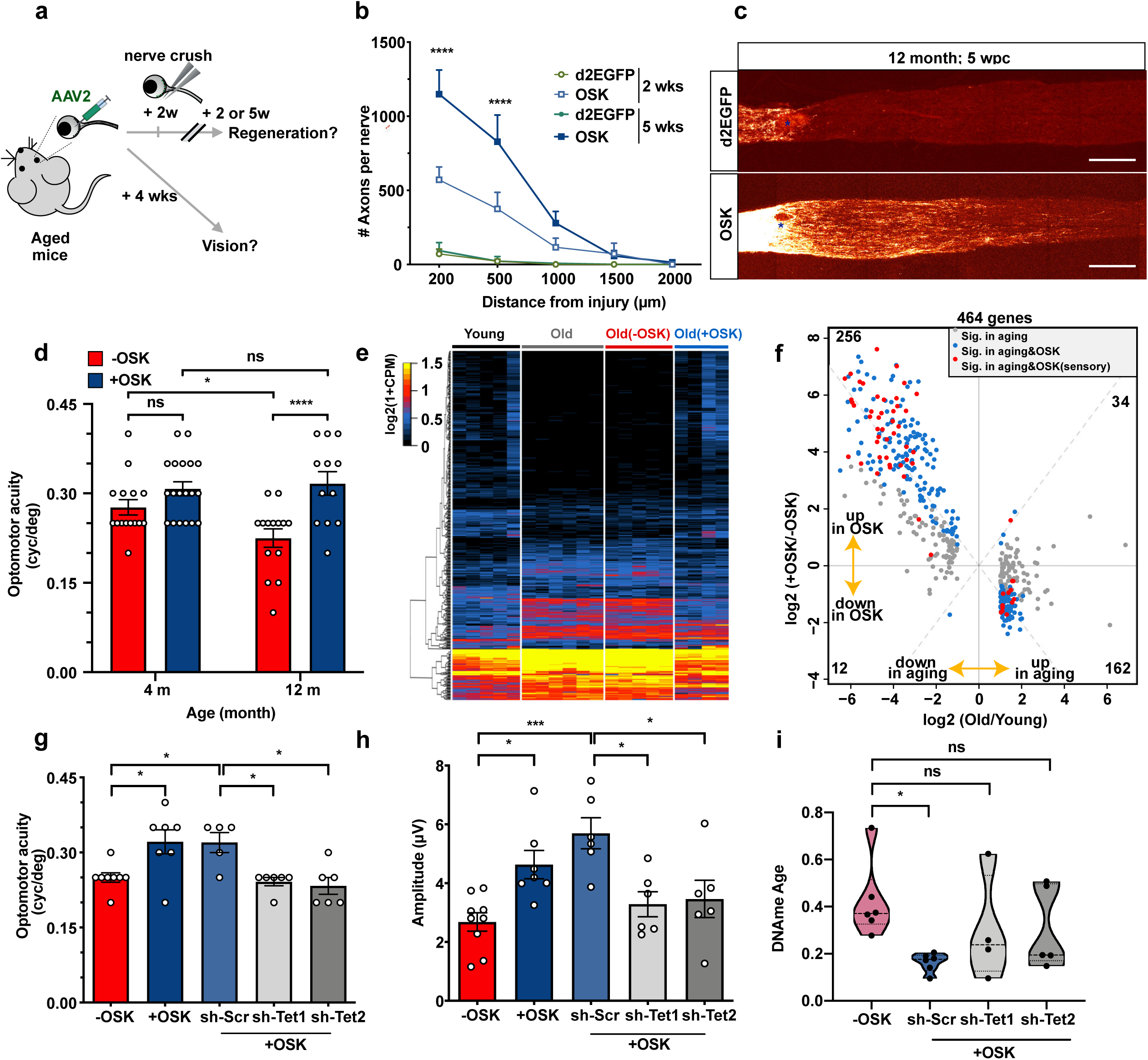
OSK AAV treatment in aged mice induces axon regeneration following optic nerve injury and restores visual function in aged mice. **a,** Experimental outline. **b** Axon regeneration in 12-month-old mice with OSK AAV or control AAV (d2EGFP) treatment 2 or 5 weeks after optic nerve crush. **c**, Representative confocal images of longitudinal sections through the optic nerve showing CTB-labeled axons after 5 weeks of OSK treatment. Scale bar = 200 µm. **d**, The spatial frequency threshold in young mice (4 months) and old mice (12 months) treated with -OSK or +OSK AAV for 4 weeks. **e**, Hierarchical clustered heatmap showing RNA-Seq expression of 464 differentially expressed genes in FACS sorted RGCs from intact young mice (5 months) or intact old mice (12 months), or old mice treated with either -OSK or +OSK AAV. -OSK: AAV-TRE-OSK; +OSK: AAV-tTA+AAV-TRE-OSK. **f,** Scatter plot of OSK-induced changes versus age-associated changes in mRNA levels. Dots represent differentially expressed genes in RGCs. **g** and **h**, Spatial frequency threshold and PERG amplitudes in old mice (12 months) treated with: (i) -OSK, (ii) +OSK, or (iii) +OSK together with either sh-Scr or sh-Tet1/sh-Tet2-mediated knockdown for 4 weeks. -OSK: AAV-rtTA+AAV-TRE-OSK; +OSK: AAV-tTA+AAV-TRE-OSK. **i**, rDNA methylation age of 12-month-old RGCs FACS isolated from retinas infected for 4 weeks with -OSK or +OSK AAV together with short-hairpin DNAs with a scrambled sequence (sh-Scr) or targeted to Tet1 or Tet2 (sh-Tet1/sh-Tet2). Gene exclusion criteria for **e** and **f**: genes with low overall expression (log2(CPM)<2), genes that did not significantly change with age (absolute log2 fold-change <1) or genes altered by the virus (differentially expressed between intact old and old treated with TRE-OSK AAV). \**P* < 0.05; \*\**P* < 0.01; \*\*\**P* < 0.001, \*\*\*\**P* < 0.0001. One-way ANOVA with Bonferroni correction in **b**, **g**, **h** and **i,** relative to first group in **i.** Two-way ANOVA with Bonferroni correction in **d.**

Glaucoma, a progressive loss of RGCs and their axons that often coincides with increased intraocular pressure, is a leading cause of age-related blindness worldwide. Although some treatments can slow down disease progression^35^, it is currently not possible to restore vision once it has been lost. Given the ability of OSK to regenerate axons after acute nerve damage, we tested whether REVIVER could restore the function of RGCs in a chronic setting like glaucoma (Fig. 3a). Elevated intraocular pressure (IOP) was induced unilaterally for 4-21 days by injection of microbeads into the anterior chamber (Fig. 3b)^36^. AAVs or PBS were then injected intravitreally and expressed at a time point when glaucomatous damage was established, with a significant decrease in RGCs and axonal density (Fig. 3a, Extended Data Fig.9a, b). Four weeks after AAV injection, OSK-treated mice presented with a significant increase in axon density compared to mice that received either PBS or AAVs with no OSK induction (-OSK). The increased axon density was equivalent to the axon density in the saline-only, non-glaucomatous mice (Fig. 3c, d) and was not associated with proliferation of RGCs (Extended Data Fig.9c).

To determine whether the increased axon density in OSK-treated mice coincided with increased vision, we tracked the visual acuity of each mouse by measuring their optomotor response (OMR) (Fig. 3e). Compared to mice that received either PBS or -OSK AAV, those that received OSK induction experienced a significant increase in visual acuity relative to the pre-treatment baseline measurement, restoring about half of vision (Fig. 3f). A readout of electrical waves generated by RGCs in response to a reversing contrast checkerboard pattern, known as pattern electroretinogram (PERG) analysis, showed that OSK treatment significantly improved RGC function relative to the pre-treatment baseline measurements and PBS or -OSK AAV treatments (Fig. 3g, h). To our knowledge, REVIVER is the first treatment to reverse vision loss in a glaucoma model.

Many treatments known to work well in young individuals often fail in older ones. For example, an approach to regenerate retinal rod photoreceptors was effective in 1 month-old mice but not in 7 month-olds^37^. Given the ability of REVIVER to regenerate axons and to restore vision after glaucomatous damage in young mice, we wondered if REVIVER might be effective in aged mice, too. Optic nerve crush injury was performed on 12 month-old mice using the same protocol as in Fig.1d according to the experimental design in Fig 4a. In aged mice, OSK AAV treatment for two weeks post-injury doubled RGC survival, similar to that observed in 1 and 3 month-old mice (Extended Data Fig. 10a). Though the axon regeneration was slightly less than young mice two weeks after injury (Fig. 4b, Extended Data Fig. 10b), OSK AAV treatment in aged mice for five weeks was similar to that observed in young mice (Fig. 4b, c). These data indicate that ageing does not significantly diminish the effectiveness of OSK AAV treatment in inducing axon regeneration following an optic nerve crush injury.

To test whether REVIVER could reverse vision loss associated with physiological ageing, 4 and 12 month-old mice received intravitreal injections of -OSK or +OSK AAV (Fig 4a). Compared to the 4 month-olds, there was a significant reduction in visual acuity and RGC function at one year of age, as measured by OMR and PERG. Strikingly, this loss was completely restored by 4-weeks of OSK expression (Fig. 4d, Extended Data Fig.10c). We did not see a restorative effect in 18 month-old mice (Extended Data Fig. 10c, d), likely due to spontaneous corneal opacity that develops at that age^38^.

Considering there was no obvious increase in RGCs and axon density in the 12m-old-mice (Extended Data Fig. 10e, f), we suspected the increased vision was due to a functional improvement, one that could be revealed by analyzing the transcriptome. FACS-purified RGCs from 12-m-old mice, either untreated or treated with -OSK or +OSK AAV, were analyzed by RNA-seq. Compared to RGCs from 5-m-old young mice, we identified 464 genes that were differentially-expressed during ageing and also not affected by empty AAV infection (Extended Data Fig. 11a and Supplemental Table 1-4). Remarkably, the vast majority (90%, 418) of the 464 age-deregulated genes were restored to youthful levels after 4 weeks of OSK expression (Fig. 4e, f). Of the 268 age-downregulated genes, 44 were genes involved in sensory perception (Fig.4f), suggesting a decline in signaling receptors or sensory function during ageing, one that can be restored by REVIVER (Extended Data Fig. 11b, c). Interestingly, 116 of these genes are yet to be characterized. Another 196 genes that were up-regulated during ageing are known or predicted to be involved in ion transport (Extended Data Fig.11d).

Consistent with the axon regeneration data, the knockdown of Tet1 or Tet2 completely blocked the ability of REVIVER to restore vision in 12 month-old mice (Fig. 4g, h). To determine if the DNA methylation clock was affected, we measured the rDNA methylation age of FACS-sorted RGCs from 12 month-old mice. Four weeks of OSK AAV expression significantly decreased DNA methylation age, and this was Tet1- and Tet2-dependent (Fig. 4i). Together, these results demonstrate that Tet-dependent *in vivo* reprogramming can restore youthful gene expression patterns, reverse the DNA methylation clock, and restore the function and regenerative capacity of a tissue as complex as the retina.

Post-mitotic neurons in the CNS are some of the first cells in the body to lose their ability to regenerate. In this study, we show that *in vivo* reprogramming of aged neurons can reverse DNA methylation age and allow them to regenerate and function as though they were young again. The requirement of the DNA demethylases Tet1 and Tet2 for this process indicates that altered DNA methylation patterns may not just a measure of age but participants in ageing. These data lead us to conclude that mammalian cells retain a set of original epigenetic information, in the same way Shannon’s observer stores information to ensure the recovery of lost information^17^. How cells are able to mark and retain youthful DNA methylation patterns, then in late adulthood OSK can instruct the removal of deleterious marks is unknown. Youthful epigenetic modifications may be resistant to removal by the Tets by the presence of a specific protein or DNA modification that inhibits the reprogramming machinery. Even in the absence of this knowledge, these data indicate that the reversal of DNA methylation age and the restoration of a youthful epigenome could be an effective strategy, not just to restore vision, but to give complex tissues the ability to recover from injury and resist age-related decline.

## Acknowledgements

We greatly thank Amy Wagers, Raul Mostoslavsky, Yang Shi, Abhirup Das, Alice Kane, Margarete Karg, Bohan Zhang, Phillip Dmitriev, Keith Booher, Emily Chen, and Shurong Hou for advice and assistance. The work was supported by the Harvard Medical School Epigenetics Seed Grant Program and Development Grant Program, The Paul F. Glenn Foundation for Medical Research, a kind gift from Edward Schulak, and NIH awards R01AG019719 (to D.A.S), R01EY026939 and R01EY021526 (to Z.H.), and R01GM065204 (to V.N.G.). We thank Boston Children’s Hospital Viral Core, which is supported by NIH5P30EY012196, and Schepen Eye Institute Core facilities, supported by NEI-P30EY003790. X.T. was supported by NASA Postdoctoral Fellowship 80NSSC19K0439; D.V. by NIH training grant T32AG023480; J.-H.Y. was partially supported by National Research Foundation of Korea (2012R1A6A3A03040476); B.R.K. partially by the St Vincent de Paul Foundation; and M.G.K. by NEI award R21EY030276. This paper is dedicated to Honghua Lu, grandfather to Y.L., who passed away bravely fighting ageing during preparation of this manuscript.

## Contributions

Y.L., X.T. and D.A.S. wrote the manuscript with input from coauthors. Y.L. and D.A.S conceived of the project with help from M.S.B.. Y.L. performed or involved in all experiments and analysis. X.T. conducted human neuron experiments. B.B., C.W., Q.Z., D.Y., S.Z., and Z.H. contributed to the optic nerve crush studies and imaging. A.K., Q.Z., D.Y., E.M.H., E.K., M.G.K., and B.R.K., contributed to the glaucoma and ageing studies. X.T., J.-H.Y., and K.H. helped with transgenic mice work. M.S.B., M.B.S., L.R., helped with systemic AAV9 experiment. M.M. and V.G. conducted mouse RGC methylation clock analysis. D.V. performed the RNA-seq analysis. N.D., and G.C. helped with plasmid constructs and AAV9 production. S.H. conducted human methylation clock analysis. K.C. helped with grant applications and project management. M.G.K., B.R.K., Z.H. and D.A.S. jointly supervised this work.

## Methods

### Mouse Lines

C57BL6/J wild type mice were purchased from Jackson Laboratory (000664) for optic nerve crush and glaucoma model experiments. For ageing experiments, females from NIA Aged Rodent Colonies (https://www.nia.nih.gov/research/dab/aged-rodent-colonies-handbook) were used. Col1a1-tetOP-OKS-mCherry/ Rosa26-M2rtTA alleles were a gift from the Hochedlinger lab (Harvard).^39^ All animal work was approved by Harvard Medical School, Boston Children’s Hospital, and Mass Eye and Ear Institutional animal care and use committees.

### Surgery

Mice were anesthetized by intraperitoneal injection of a mixture of ketamine (100 mg/kg; Ketaset; Fort Dodge Animal Health, Fort Dodge, IA) and xylazine (9 mg/kg; TranquiVed; Vedco, Inc., St. Joseph, MO) supplemented by topical application of proparacaine to the ocular surface (0.5%; Bausch & Lomb, Tampa, FL). All animal procedures were approved by the IACUC of the respective institutions and according to appropriate animal welfare regulations.

### Production of Adeno associated viruses (AAVs)

Vectors of AAV-TRE-OSK were made by cloning mouse Oct4, Sox2 and Klf4 cDNA into an AAV plasmid consisting of the Tet Response Element (TRE3G promoter) and SV40 element. The other vectors were using similar strategy or directly chemically synthesized. All pAAVs, as listed (Supplemental Table 5), were then packaged into AAVs of serotype 2/2 or 2/9 (titers: > 5×10^12^ genome copies/ml). AAVs were produced by Boston Children’s Hospital Viral Core.

### Systemic delivery of AAV9 to internal organs

Expression in internal organs was achieved through retro-orbital injection of AAV9 (3×10^11^ TRE-OSK plus 7×10^11^ UBC-rtTA). To induce OSK expression, doxycycline (1 mg/ml; MP biochemicals) was given in drinking water continuously, 3 weeks post-AAV injection.

### Cell culture and differentiation

Ear fibroblasts (EFs) were isolated from Reprogramming 4F (Jackson Laboratory 011011) or 3F (Hochedlinger lab, Harvard) mice and cultured at 37°C in DMEM (Invitrogen) containing Gluta-MAX, non-essential amino acids, and 10% fetal bovine serum (FBS). EFs of transgenic OSKM and OSK mice were passaged to P3 and treated with doxycycline (2 mg/ml) for the indicated time periods in the culture medium. SH-SY5Y neuroblastoma cells were obtained from the American Tissue Culture Collection (ATCC, CRL-2266) and maintained according to ATCC recommendations. Cells were cultured in a 1:1 mixture of Eagle’s Minimum Essential Medium (EMEM, ATCC, 30-2003) and F12 medium (ThermoFisher Scientific, 11765054) supplemented with 10% fetal bovine serum (FBS, Sigma, F0926) and 1X penicillin/streptomycin (ThermoFisher Scientific, 15140122). Cells were cultured at 37°C with 5% CO_2_ and 3% O_2_. Cells were passaged at ∼80% confluency. SH-SY5Y cells were differentiated into neurons as previously described^40^ with modifications. Briefly, 1 day after plating, cell differentiation was induced for 3 days using EMEM/F12 medium (1:1) containing 2.5% FBS, 1× penicillin/streptomycin, and 10 µM all-trans retinoic acid (ATRA, Stemcell Technologies, 72264) (Differentiation Medium 1), followed by a 3 day incubation in EMEM/F12 (1:1) containing 1% FBS, 1 × penicillin/streptomycin, and 10 µM ATRA (Differentiation Medium 2). Cells were then split into 35 mm cell culture plates coated with poly-D-lysine (ThermoFisher Scientific, A3890401). A day after splitting, neurons were matured in serum-free neurobasal/B27 plus culture medium (ThermoFisher Scientific, A3653401) containing 1 × Glutamax (ThermoFisher Scientific, 35050061), 1 × penicillin/streptomycin, and 50 ng/ml BDNF (Alomone labs) (Differentiation Medium 3) for at least 5 days.

### Neurite regeneration assay

Differentiated SH-SY5Y cells were transduced with AAV.DJ vectors at 10^6^ genome copies/cell. Five days after transduction, vincristine (100 nM; Sigma, V8879) was added for 24 hrs to induce neurite degeneration. Neurons were then washed twice in PBS and fresh Differentiation medium 3 was added back to the plates. Neurite outgrowth was monitored for 2-3 weeks by taking phase-contrast images at 100x magnification every 3-4 days. Neurite area was quantified using Image J.

### Cell cycle analysis

Cells were harvested and fixed with 70% cold ethanol for 16 hrs at 4°C. After fixation, cells were washed twice with PBS and incubated with PBS containing 50 μg/mL propidium iodide (Biotium, 40017) and 100 μg/mL RNase A (Omega) for 1 hr at room temperature. PI-stained samples were analyzed on a BD LSR II analyzer and only single cells were gated for analysis. Cell cycle profiles were analyzed using FCS Express 6 (De Novo Software).

### Human neuron methylation and epigenetic clock analyses

DNA was extracted using the Zymo Quick DNA mini-prep plus kit (D4069) and DNA methylation levels were measured on Illumina 850 EPIC arrays. The Illumina BeadChip (EPIC) measured bisulfite-conversion-based, single-CpG resolution DNAm levels at different CpG sites in the human genome. Data were generated via the standard protocol of Illumina methylation assays, which quantifies methylation levels by the β value using the ratio of intensities between methylated and un-methylated alleles. Specifically, the β value was calculated from the intensity of the methylated (M corresponding to signal A) and un-methylated (U corresponding to signal B) alleles, as the ratio of fluorescent signals β = Max(M,0)/(Max(M,0)+ Max(U,0)+100). Thus, β values ranged from 0 (completely un-methylated) to 1 (completely methylated). “Noob” normalization was implemented using the “minfi” R package^41, 42^. The mathematical algorithm and available software underlying the skin & blood clock for *in vitro* studies (based on 391 CpGs) was previously published^34^.

### AAV2 Virus Intravitreal Injection

Adult animals were anesthetized with ketamine/xylazine (100/10 mg/kg) and then AAV (1-3 µl) was injected intravitreally, just posterior to the limbus with a fine glass pipette attached to the Hamilton syringe using plastic tubing. In elevated IOP model, mice received a 1µl intravitreal injection between 3-4 weeks following microbead injection. The injected volume of AAV-sh-RNA is 1/5th the volume of other AAVs.

### Optic Nerve Crush

Two weeks after intravitreal AAV injection, the optic nerve was accessed intraorbitally and crushed in anesthetized animals using a pair of Dumont #5 forceps (FST). Alexa-conjugated cholera toxin beta subunit (CTB-555, 1 mg/ml; 1-2 µl) injection was performed 2-3 days before euthanasia to trace regenerating RGC axons. More detailed surgical methods were described previously^24^.

### *In Vivo* Doxycycline Induction or suppression

Induction of Tet-On or suppression of Tet-Off AAV2 systems in the retina was performed by administration of doxycycline (2 mg/ml) (Sigma) in the drinking water. Induction of Tet-On AAV9 system systemically was performed by administration of doxycycline (1 mg/ml) (USP grade, MP Biomedicals 0219895505) in the drinking water.

### Axon Regeneration Quantification

The number of regenerating axons in the optic nerve was estimated by counting the number of CTB-labeled axons at different distances from the crush site as described previously^24^.

### Whole-Mount Optic Nerve Preparation

Optic nerves and the connecting chiasm were dehydrated in methanol for 5 min, then incubated overnight with Visikol® HISTO-1™. Next day nerves were transferred to Visikol® HISTO-2™ and then incubated for 3 hrs. Finally, optic nerves and connecting chiasm were mounted with Visikol® HISTO-2™.

### Immunofluorescence

Whole-mount retinas were blocked with horse serum 4°C overnight then incubated at 4°C for 3 days with primary antibodies: Mouse anti-Oct4 (1:100, BD bioscience, 611203), Rabbit anti-Sox2 (1:100, Cell signaling, 14962), Goat anti-Klf4 (1:100, R&D system, AF3158), Rabbit anti-phosphorylated S6 Ser235/236 (1:100, Cell Signaling 4857), Rabbit anti-Brn3a (1:200, EMD Millipore, MAB1585) and Guinea pig anti-RBPMS (1:400, Raygene custom order A008712 to peptide GGKAEKENTPSEANLQEEEVRC) diluted in PBS, BSA (3%) Triton X-100 (0.5%). Then, tissues were incubated at 4°C overnight with appropriate Alexa Fluor-conjugated secondary antibodies (Alexa 405, 488, 567, 674; Invitrogen) diluted with the same blocking solution as the primary antibodies, generally used at 1:400 final dilution. Frozen sections were stained overnight with primary antibodies at 4°C and then secondary antibodies at room temperature for 2 h. Sections or whole-mount retinas were mounted with Vectashield Antifade Mounting Medium.

### Western Blot

SDS-PAGE and western blot analysis was performed according to standard procedures and detected with an ECL detection kit. Antibodies used: Rabbit anti-Sox2 (1:100, EMD Millipore, AB5603), Mouse anti-Klf4 (1:1000, ReproCell, 09-0021), Rabbit anti-p-S6 (S240/244) (1:1000, CST, 2215), Mouse anti-S6 (1:1000, CST, 2317), Mouse anti-β-Tubulin (1:1000, Sigma-Aldrich, 05-661), Mouse anti-β-Actin−Peroxidase antibody (1:20,000, Sigma-Aldrich, A3854).

### RGC Survival and Phospho-S6 Signal

RBPMS-positive cells in the ganglion layer were stained with an anti-RBPMs antibody (1:400, Raygene custom order A008712 to peptide GGKAEKENTPSEANLQEEEVRC) and a total of four 10X fields per retina, one in each quadrant, were enumerated. The average number per field was determined and the percentages of viable RGCs were obtained by comparing the values determined from the uninjured contralateral retinas. Phospho-S6 (1:100, Cell Signaling 4857) staining was performed under the same conditions and the densities of phopsho-S6-positive RGCs were obtained by comparing the value to uninjured contralateral retinas.

### Microbead-induced mouse model of elevated IOP

Elevation of IOP was induced unilaterally by injection of polystyrene microbeads (FluoSpheres; Invitrogen, Carlsbad, CA; 15-μm diameter) to the anterior chamber of the right eye of each animal under a surgical microscope, as previously reported^36^. Briefly, microbeads were prepared at a concentration of 5.0 × 10^6^ beads/mL in sterile physiologic saline. A 2 μL volume was injected into the anterior chamber through a trans-corneal incision using a sharp glass micropipette connected to a Hamilton syringe (World Precision Instruments Inc., Sarasota, FL) followed by an air bubble to prevent leakage. Any mice that developed signs of inflammation (clouding or an edematous cornea) were excluded.

### IOP (Intraocular pressure) measurements

IOPs were measured with a rebound TonoLab tonometer (Colonial Medical Supply, Espoo, Finland), as previously described^36, 43^. Mice were anesthetized by 3% isoflurane in 100% oxygen (induction) followed by 1.5% isoflurane in 100% oxygen (maintenance) delivered with a precision vaporizer. IOP measurement was initiated within 2-3 min after the loss of a toe or tail pinch reflex. Anesthetized mice were placed on a platform and the tip of the pressure sensor was placed approximately 1/8 inch from the central cornea. Average IOP was displayed automatically after 6 measurements after elimination of the highest and lowest values. The machine-generated mean was considered as one reading, and six readings were obtained for each eye. All IOPs were taken at the same time of day (between 10:00 and 12:00 hrs) due to the variation of IOP throughout the day.

### Optomotor Response

Visual acuity of mice was measured using an optomotor reflex-based spatial frequency threshold test^44, 45^. Mice were able to freely move and were placed on a pedestal located in the center of an area formed by four computer monitors arranged in a quadrangle. The monitors displayed a moving vertical black and white sinusoidal grating pattern. A blinded observer, unable to see the direction of the moving bars, monitored the tracking behavior of the mouse. Tracking was considered positive when there was a movement of the head (motor response) to the direction of the bars or rotation of the body in the direction concordant with the stimulus. Each eye would be tested separately depending on the direction of rotation of the grating. The staircase method was used to determine the spatial frequency start from 0.15 to 0.40 cycles/deg, with intervals of 0.05 cycles/deg. Rotation speed (12°/s) and contrast (100%) were kept constant. Responses were measured before and after treatment by individuals blinded to the group of the animal and the treatment.

### Pattern Electroretinogram (PERG)

Mice were anesthetized with ketamine/xylazine (100mg/kg and 20mg/kg) and the pupils dilated with one drop of 1% tropicamide ophthalmic solution. The mice kept under dim red light throughout the procedure on a built-in warming plate (Celeris, Full-Field and Pattern Stimulation for the rodent model) to maintain body temperature at 37°C. A black and white reversing checkerboard pattern with a check size of 1° was displayed on light guide electrode-stimulators placed directly on the ocular surface of both eyes and centered with the pupil. The visual stimuli were presented at 98% contrast and constant mean luminance of 50 cd/m^2^, spatial frequency: 0.05 cyc/deg; temporal frequency: 1Hz. A total of 300 complete contrast reversals of PERG were repeated twice in each eye and the 600 cycles were segmented and averaged and recorded. The averaged PERGs were analyzed to evaluate the peak to trough N1 to P1 (positive wave) amplitude.

### Quantification of optic nerve axons

For quantification of axons, optic nerves were dissected and fixed overnight in Karnovsky’s reagent (50% in phosphate buffer). Semi-thin cross-sections of the nerve were taken at 1.0 mm posterior to the globe and stained with 1% p-phenylenediamine (PPD) for evaluation by light microscopy. Optic nerve cross sections were imaged at 60x magnification using a Nikon microscope (Eclipse E800, Nikon, Japan) with the DPController software (Olympus, Japan) and 6-8 non-overlapping photomicrographs were taken to cover the entire area of each optic nerve cross-section. Using ImageJ (Version 2.0.0-rc-65/1.51u), a 100 μM x 100 μM square was placed on each 60x image and all axons within the square (0.01mm^2^) were counted using the threshold and analyze particles function in image J as previously described^36, 43, 44^. Damaged axons stain darkly with PPD and are not counted. The average axon counts in the 6-8 images were used to calculate the axon density/mm^2^ of optic nerve. Individuals performing axon counts were blinded to the experimental groups.

### Quantification of retinal ganglion cells in glaucoma model

For ganglion cell counting, images of whole mount retinas were acquired using a 63x oil immersion objective of the Leica TCS SP5 confocal microscope (Leica Microsystems). The retinal whole mount was divided into four quadrants and two to four images (248.53μm by 248.53μm in size) were taken from the midperipheral and peripheral regions of each quadrant, for a total of twelve to sixteen images per retina. The images were obtained as z-stacks (0.5μm) and all Brn3a positive cells in the ganglion cell layer of each image were counted manually as previously described^44^. Briefly, RGCs were counted using the “Cell Counter” plugin (http://fiji.sc/Cell_Counter) in Fiji is Just ImageJ software (ImageJ Fiji, version 2.0.0-rc-69/1.52n). Each image was loaded into Fiji and a color counter type was chosen to mark all Brn3a stained RGCs within each image (0.025mm ^2^). The average number of RGCs in the 12 to sixteen images were used to calculate the RGC density per square millimeter of retina. Two individuals blinded to the experimental groups performed all RGC counts.

### RGC Enrichment

Retinas were fresh dissected and dissociated in AMES media using papain, then triturated carefully and stained with Thy1.2-PE-Cy7 anti-body (Invitrogen 25-0902-81) and Calcine Blue live-dead cell stain, then flow sorted on a BD FACS Aria using an 130µm nozzle to collect over 10,000 Thy1.2+ and Calcine blue+ cells (1-2% of total events). Frozen cells were sent to GENEWIZ, LLC (South Plainfield, NJ, USA) for RNA extraction and ultra-low input RNA-seq, or to Zymo research (Irving, CA) for DNA extraction and genome-wide reduced representation bisulfite sequencing (RRBS).

### Classic RRBS Library preparation

DNA was extracted using Quick-DNA Plus Kit Microprep Kit. 2-10 ng of starting input genomic DNA was digested with 30 units of *Msp*I (NEB). Fragments were ligated to pre-annealed adapters containing 5’-methyl-cytosine instead of cytosine according to Illumina’s specified guidelines. Adaptor-ligated fragments ≥50 bp in size were recovered using the DNA Clean & ConcentratorTM-5 (Cat#: D4003). The fragments were then bisulfite-treated using the EZ DNA Methylation-LightningTM Kit (Cat#: D5030). Preparative-scale PCR products were purified with DNA Clean & ConcentratorTM-5 (Cat#: D4003) for sequencing on an Illumina HiSeq using 2×125 bp PE.

### DNA methylation age analysis of mouse RGC

Reads were filtered using trim galore v0.4.1 and mapped to the mouse genome GRCm38 using Bismark v0.15.0. Methylation counts on both positions of each CpG site were combined. Only CpG sites covered in all samples were considered for analysis. This resulted in total of 708156 sites. For the rDNA methylation clock reads were mapped to BK000964 and the coordinates were adjusted accordingly^26^. 70/72 sites were covered for rDNA clock, compared to 102/435 sites of whole lifespan multi-tissue clock^27^, or 248/582 and 77,342/ 193,651 sites (ridge) of two entire lifespan multi-tissue clocks^28^.

### Total RNA extraction and Sample QC

Total RNA was extracted following the Trizol Reagent User Guide (Thermo Fisher Scientific). 1 µl of 10 mg/ml Glycogen was added to the supernatant to increase RNA recovery. RNA was quantified using Qubit 2.0 Fluorometer (Life Technologies, Carlsbad, CA, USA) and RNA integrity was determined using TapeStation (Agilent Technologies, Palo Alto, CA, USA).

### Ultra-low input RNA library preparation and multiplexing

RNA samples were quantified using Qubit 2.0 Fluorometer (Life Technologies, Carlsbad, CA, USA) and RNA integrity was ascertained using a 2100 TapeStation (Agilent Technologies, Palo Alto, CA, USA). RNA library preparations, sequencing reactions, and initial bioinformatics analysis were conducted at Genewiz (South Plainfield, NJ, USA). A SMART-Seq v4 Ultra Low Input Kit for Sequencing was used for full-length cDNA synthesis and amplification (Clontech, Mountain View, CA), and Illumina Nextera XT library was used for sequencing library preparation. Briefly, cDNA was fragmented and adaptors were added using Transposase, followed by limited-cycle PCR to enrich and add an index to the cDNA fragments. The final library was assessed by a Qubit 2.0 Fluorometer and Agilent TapeStation.

### Sequencing 2×150bp PE

The sequencing libraries were multiplexed and clustered on two lanes of a flowcell. After clustering, the flowcell were loaded on the Illumina HiSeq instrument according to manufacturer’s instructions. Samples were sequenced using a 2×150 Paired End (PE) configuration. Image analysis and base calling were conducted by the HiSeq Control Software (HCS) on the HiSeq instrument. Raw sequence data (.bcl files) generated from Illumina HiSeq was converted into fastq files and de-multiplexed using Illumina bcl2fastq v2.17 program. One mis-match was allowed for index sequence identification.

### RNA-seq analysis

Paired-end reads were aligned with hisat2 v2.1.0^46^ to the Ensembl GRCm38 primary assembly using splice junctions from the Ensembl release 84 annotation. Paired read counts were quantified using featureCounts v1.6.4^47^ using reads with a MAPQ >=20. Differentially-expressed genes for each pairwise comparison were identified with edgeR v3.26^48^, testing only genes with at least 0.1 counts-per-million (CPM) in at least three samples. Gene ontology analysis of differentially-expressed genes was performed with AmiGO v2.5.12^49–51^. Age-associated sensory perception genes were extracted from the mouse Sensory Perception (GO:0007600) category the Gene Ontology database, including genes that were differentially expressed (q<=0.05) in 12 versus 5 month-old mice, excluding genes that were induced by the Control virus alone (q<=0.1).

## Data availability

The data that support the findings of this study are available from the corresponding author upon reasonable request.

---

**Extended Data Fig. 1.**
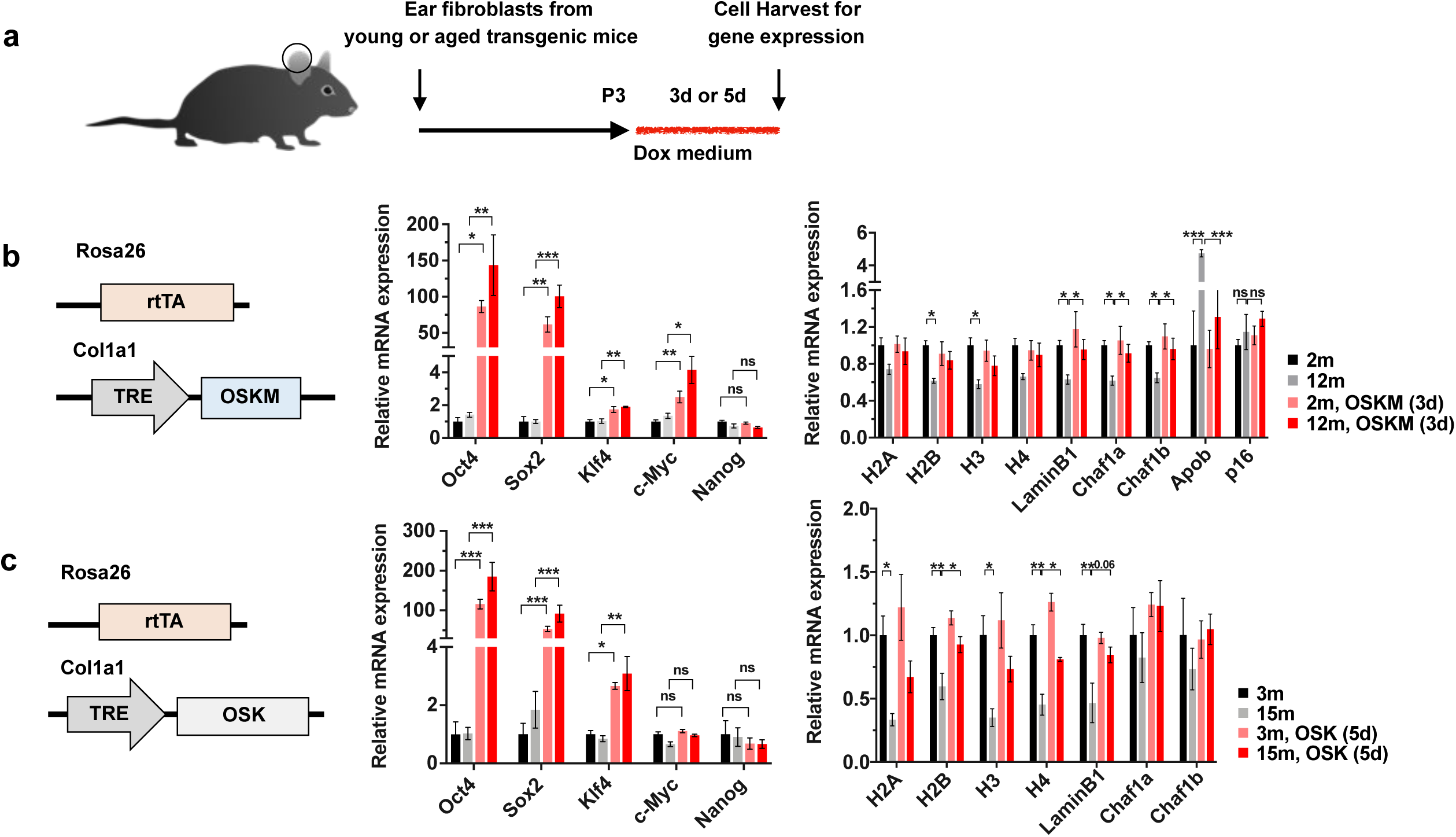
Exploration of the effect of OSK (no Myc) on ageing cells. **a**, Experimental outline for testing the rejuvenation effects of OSKM and OSK expression in young and old transgenic mouse fibroblasts. **b** and **c**, OSKM and OSK expression rescues age-associated transcriptional changes without inducing Nanog expression. Sequences of qPCR primers are in Supplementary Table 6. *P < 0.05; **P < 0.01; ***P < 0.001, ****P < 0.0001. Two-tailed Student’s t test.

**Extended Data Fig. 2.**
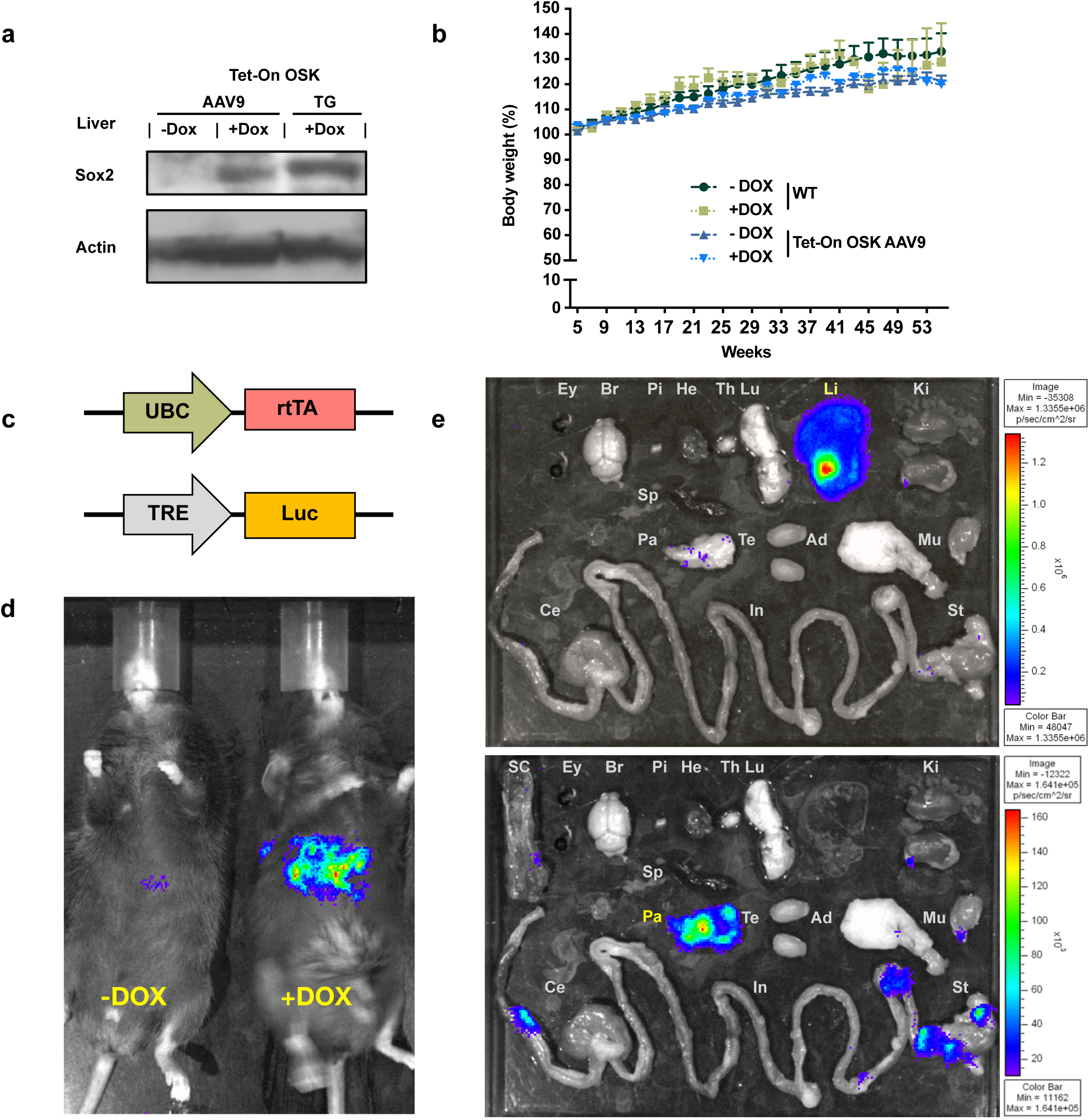
Safety of OSK AAV. **a**. Sox2 expression in the liver of WT mice post-intravenous delivery of OSK-AAV9 and OSK transgenic (TG) mice. **b**. Body weight of WT mice and AAV-mediated OSK-expressing mice (1.0×10^12 gene copies total) with or without doxycycline in the following 9 months after first 4 weeks monitoring in Fig.1b (n=5,3,6,4 respectively). **c**. AAV-UBC-rtTA and AAV-TRE-Luc vectors used for measuring tissue distribution. **d**. Luciferase imaging of WT mice at 2 months after retro-orbital injections of AAV9-UBC-rtTA and AAV9-TRE-Luc (1.0×10^12 gene copies total). Doxycycline was delivered in drinking water (1 mg/mL) for 7 days to the mouse shown on the right. **e**. Luciferase imaging of eye (Ey), brain (Br), pituitary gland (Pi), heart (He), thymus (Th), lung (Lu), liver (Li), kidney (Ki), spleen (Sp), pancreas (Pa), testis (Te), adipose (Ad), muscle (Mu), spinal cord (SC), stomach (St), small intestine (In), and cecum (Ce) 2 months after retro-orbital injection of AAV9-UBC-rtTA and AAV9-TRE-Luc followed by treatment with doxycycline for 7 days. The luciferase signal is primarily in liver. Imaging the same tissues with a longer exposure time (lower panel) revealed lower levels of luciferase signal in pancreas (liver was removed).

**Extended Data Fig.3.**
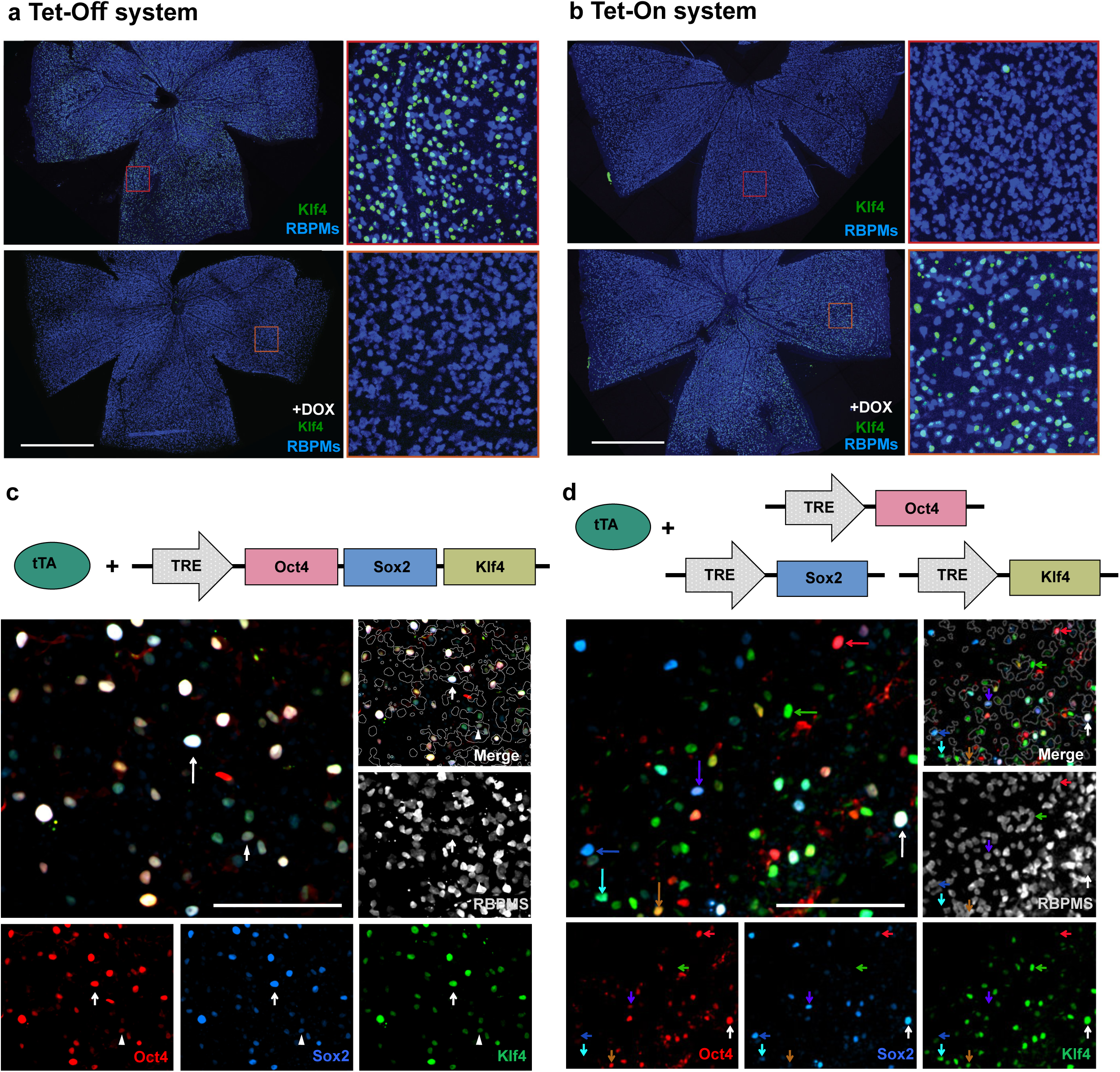
Characterization of inducible polycistronic AAV system. **a, b** Whole-mounted retina display of RBPMS and Klf4 immunofluorescence, showing that expression from the AAV2 Tet-Off system can be turned off by doxycycline in drinking water (2mg/mL 3 days), and expression from the AAV2 Tet-On system can be turned on by doxycycline drinking water (2mg/mL 2 days). Scale bars = 1 mm. **c**, Immunofluorescence analysis of the whole-mounted retina transduced with a polycistronic AAV vector expressing Oct4, Sox2, and Klf4 in the same cell. White arrows point at triple positive cells. Scale bars represent 100 µm. **d**, Immunofluorescence analysis of the whole-mounted retina transduced with AAVs separately encoding Oct4, Sox2, and Klf4. Red, blue and green arrows point at single-positive cells, white arrow point at triple positive cell, other arrows point at double positive cells. Scale bars = 100 µm.

**Extended Data Fig. 4.**
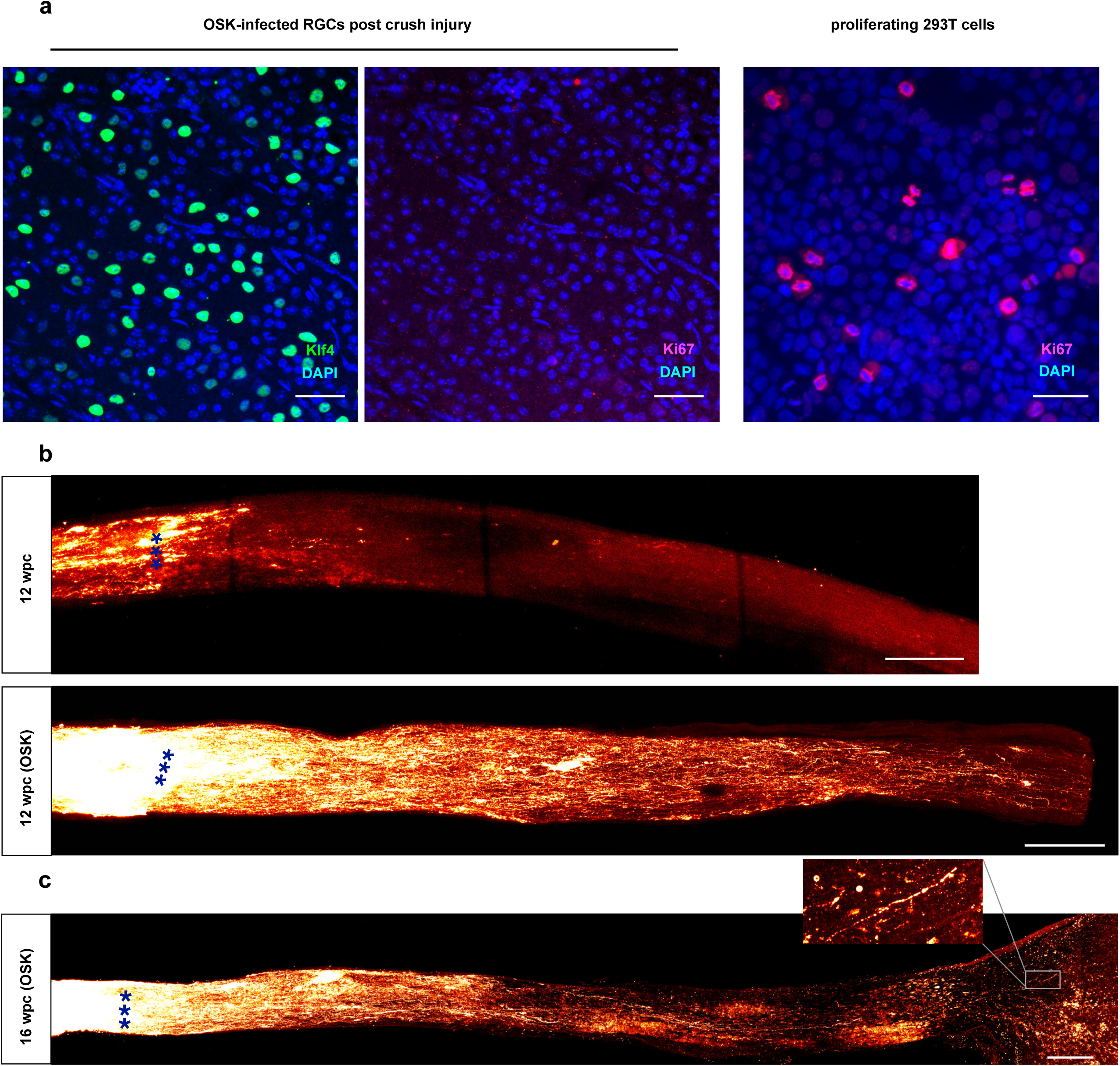
OSK induces long-term axon regeneration post-injury without RGC proliferation. **a**, Retina whole mount staining showing absence of the proliferation marker Ki67 (left). OSK infected RGCs have none while proliferating 293T cells have a signal (right). Scale bars = 100 µm. **b,** Imaging of optic nerves showing regenerating axons with or without OSK AAV treatment 12 weeks post crush (wpc). Scale bars = 200 µm. **c,** Whole nerve imaging showing CTB-labeled regenerative axons at 16 weeks post crush (wpc) in wild-type mice with intravitreal injection of AAV2-tTA and TRE-OSK. Scale bars = 200 µm.

**Extended Data Fig. 5.**
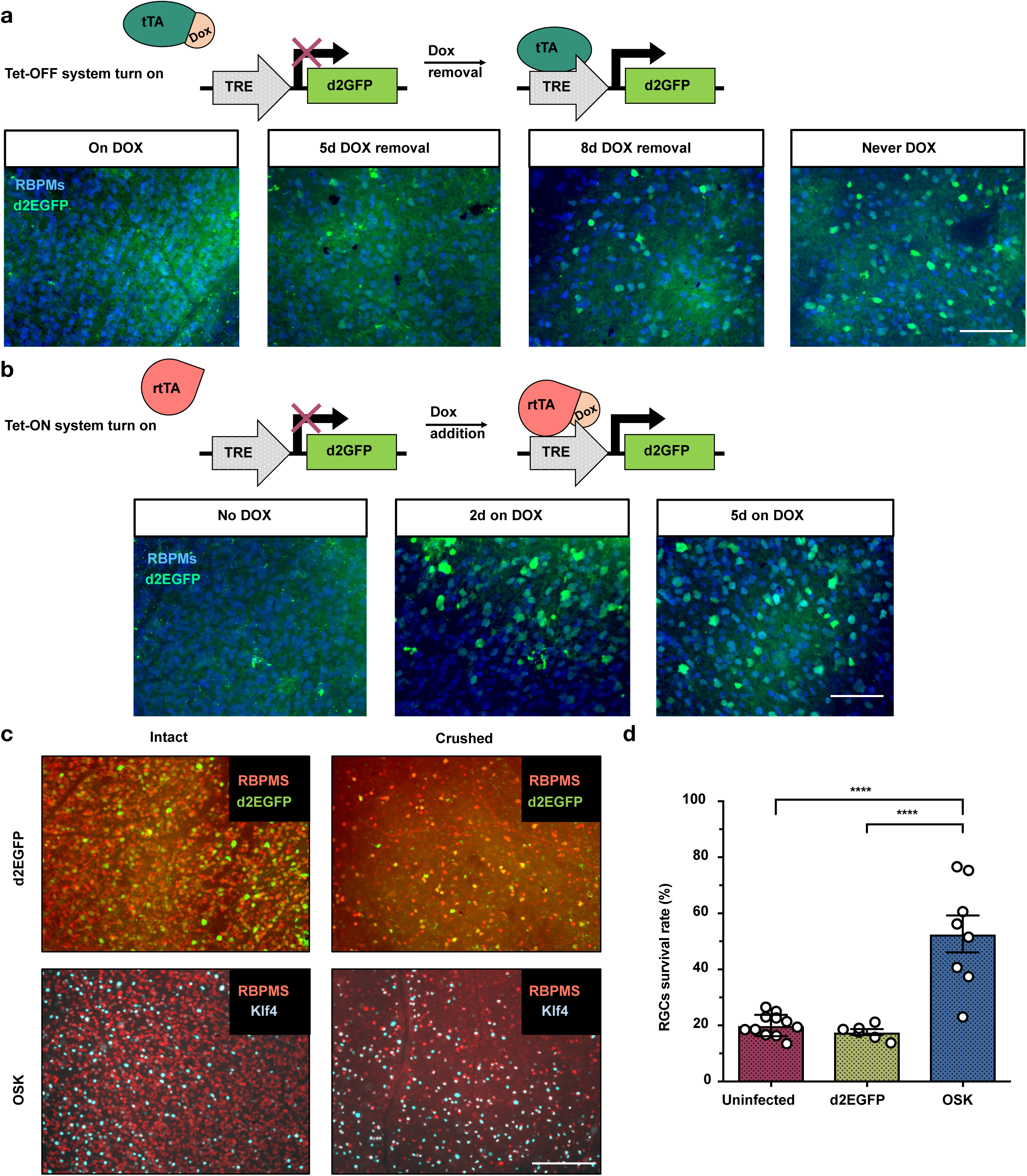
The Tet-On system has better turn on rate and OSK transduced RGCs have higher survival rate than the Tet-Off system. **a,** Representative images showing the d2EGFP expression in retina from Tet-Off AAV system with different doxycycline treatment (2 mg/mL). When pre-treated with doxycycline to suppress expression (on DOX), GFP showed up sparsely after doxycycline withdrawal for 8 days, much weaker compared to peak expression (Never DOX). **b,** Representative images showing the d2EGFP in retina from Tet-On AAV system. No GFP expression was observed in the absence of doxycycline. GFP expression reached a peak level 2 days after doxycycline induction and remained at a similar level after 5 days of induction. **c,** Representative Immunofluorescence image of GFP-positive or KLF4-positive RGCs in intact and crushed samples. **d,** Quantification of GFP- or KLF4-positive cells indicating higher survival rate of OSK expressing RGCs after crush. ****, P<0.0001, one-way ANOVA with Bonferroni correction. Scale bars = 200 µm in **a**, **b** and **c**.

**Extended Data Fig. 6.**
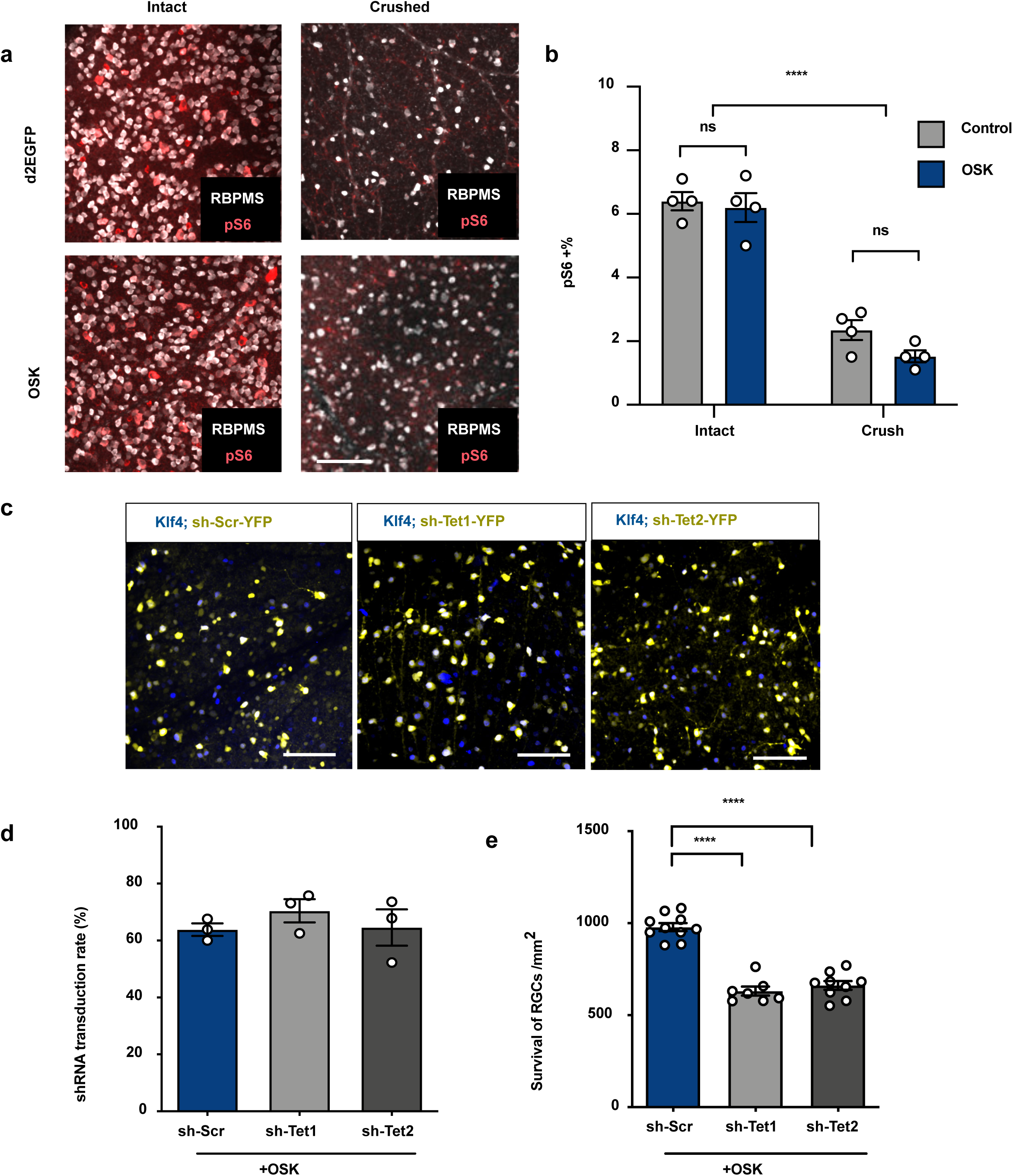
Identification of epigenetic mechanism underlying OSK effect. **a**, Representative images of retinal whole mounts transduced with d2EGFP- or OSK-encoding AAV2 in the presence or absence of crush injury. Retinal whole mounts were immunostained for the RGC marker RBPMS and mTOR activation marker pS6. **b,** Quantification of pS6-positive RGC percentile in intact and crushed samples. **c**, Representative images of retinal whole mounts transduced with TRE-OSK and tTA AAV2 in the combination with sh-Scr or sh-Tet1 or sh-Tet2 YFP AAV2 at titer ratio 5:5:1. Retinal whole mounts were immunostained for Klf4. **d**, Quantification of transduction rate of shRNA-YFP AAV in OSK expressing RGCs. **e**, Quantification of RGC survival in retinas co-transduced with AAV2 vectors encoding polycistronic OSK and tTA, in combination with sh-Scr, sh-Tet1 or sh-Tet2 YFP. Scale bars = 100 µm in **a** and **c**. *P < 0.05; **P < 0.01; ***P < 0.001, ****P < 0.0001. Two-Way ANOVA in **b**, One-way ANOVA in **e**.

**Extended Data Fig. 7.**
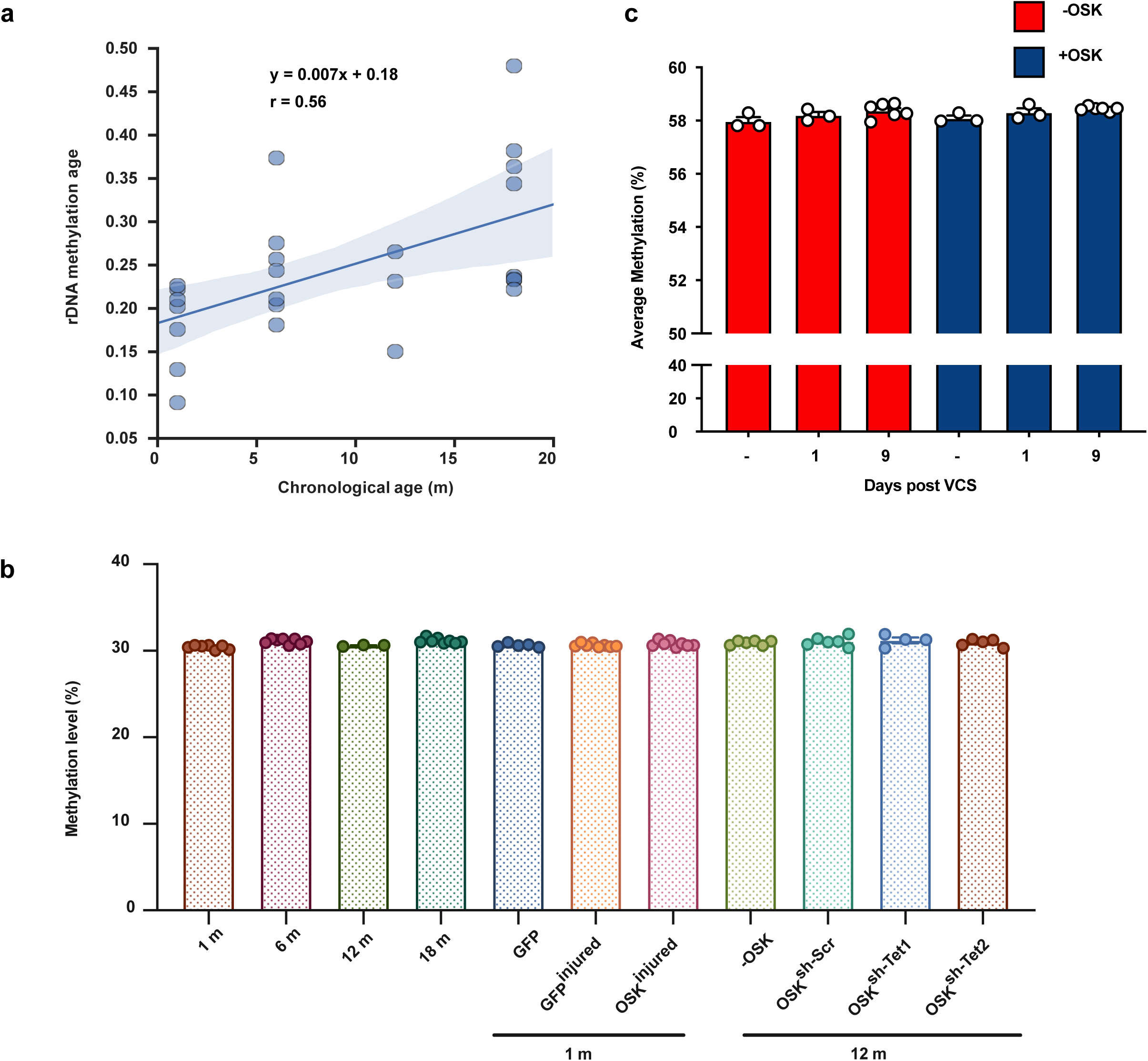
Methylation clock analysis of mouse RGCs and human neurons. **a**, Correlation between rDNA methylation age and chronological age of sorted mouse RGCs; **b**, Average DNA methylation levels of RGCs from different ages and treatments, based on 708,156 shared sites from RRBS of all samples (combined strands); **c**, Average DNA methylation levels of human neurons treated with OSK before treatment with vincristine (VCS) (–) or days post-VCS damage (1 and 9), among 850,000 probes from EPIC array.

**Extended Data Fig. 8.**
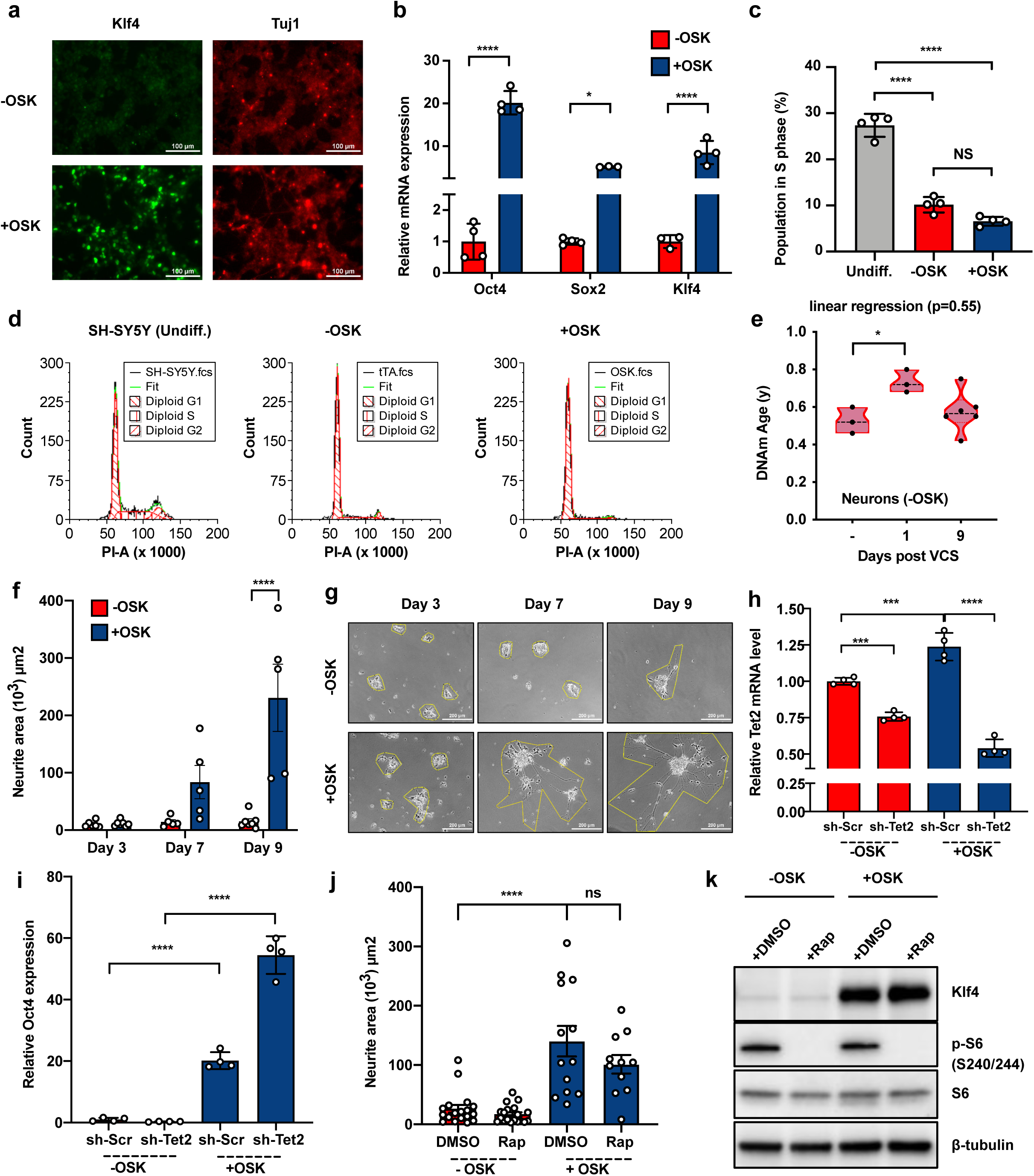
OSK induces axon regeneration of human neurons via DNA demethylases not mTOR. **a**, Immunofluorescence of differentiated human neurons with transduction of AAV-DJ vectors. -OSK: AAV-tTA; +OSK: AAV-tTA+AAV-TRE-OSK. **b,** mRNA levels of Oct4, Sox2 and Klf4 in human neurons transduced with AAV-DJ vectors. **c,** Percentage of cells in S phase. **d,** FACS profiles of G1, S, and G2 phases in undifferentiated cells and differentiated cells transduced with AAV-DJ vectors. **e**, DNA methylation age of human neurons before vincristine (VCS) damage (Day -) or 1 and 9 days post-damage in the absence of OSK expression, estimated using a skin or a blood cell clock. **f,** Quantification of neurite area at different time points after vincristine damage. **g,** Representative images and neurite area of human neurons after vincristine damage with or without OSK expression. **h** and **i**, human Tet2 mRNA level and mouse Oct4 mRNA level with sh-Scr or sh-Tet2 AAV in human neurons in the absence or presence of OSK expression. **j,** The effect of mTOR inhibition on axon regeneration of differentiated neurons with or without OSK expression. **k,** Phosphorylation level of S6 in human neurons with rapamycin treatment (10 nM) for 5 days. *p < 0.05, **p < 0.01, **** p < 0.0001, one-way ANOVA with Bonferroni’s multiple comparison test in **b**, **c**, **e**, **f**, **h**, **i**, **j**. The linear regression p value in **e** refers to DNAme Age decreasing with time.

**Extended Data Fig.9.**
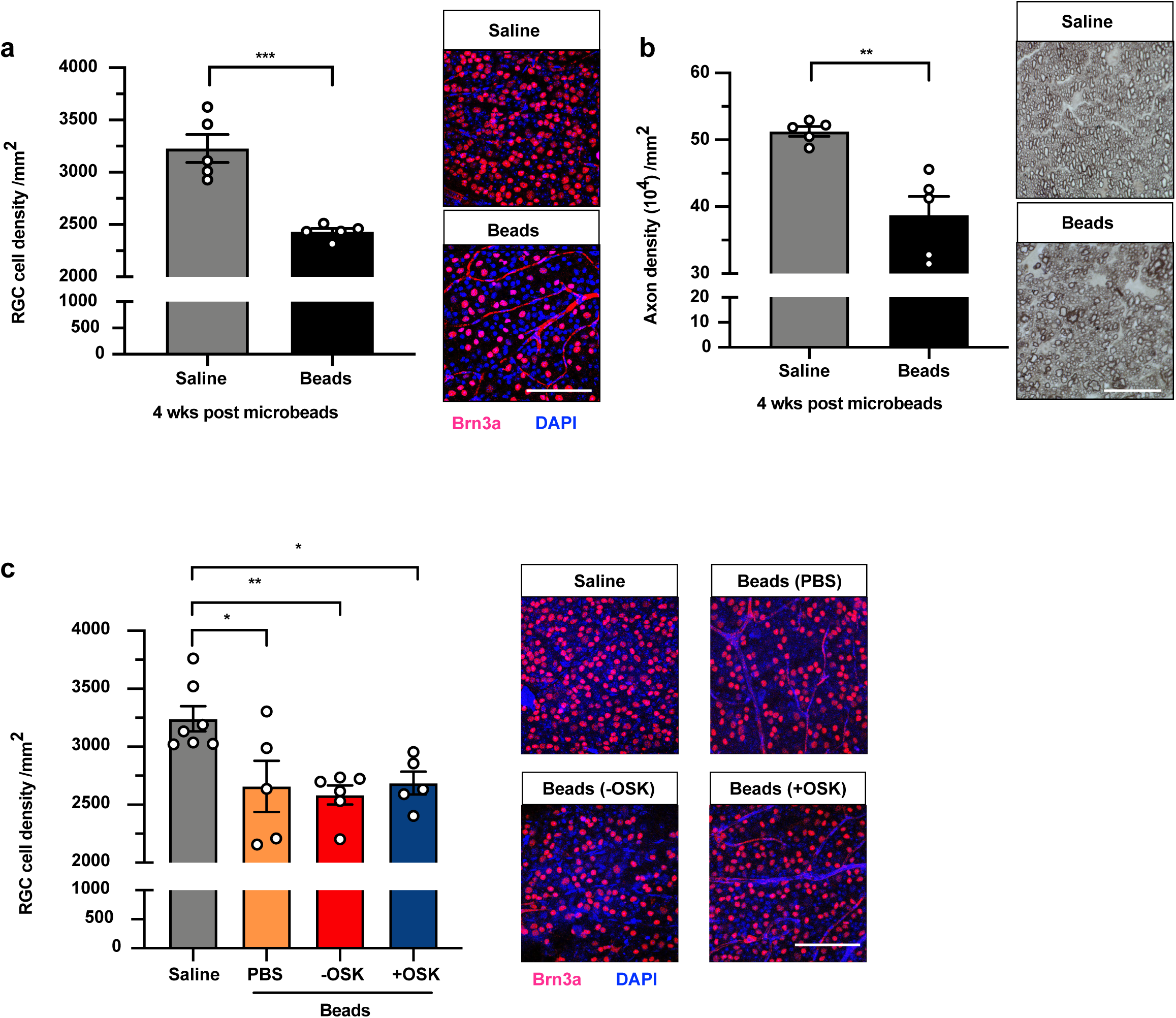
Effect of OSK in Microbead-induced glaucoma model. **a**, Quantification of RGCs and representative confocal microscopic images from retinal flat-mounts stained with anti-Brn3a (red), an RGC-specific marker, and DAPI (blue), a nuclear stain, 4 weeks after microbead or saline injection (8 weeks post-saline or microbeads). Scale bar = 100 µm. **b**, Quantification of healthy axons of optic nerve and representative photomicrographs of PPD-stained optic nerve cross-sections, 4 weeks after microbead or saline injection. Scale bars = 25 µm. **c,** Quantification of RGCs and representative confocal microscopic images 4 weeks post-PBS or AAV injection (8 weeks after a saline or microbead injection). *p < 0.05, **p < 0.01, *** p < 0.001, Student’s t-test in **a** and **b**. one-way ANOVA with Bonferroni’s multiple comparison test in **c**.

**Extended Data Fig. 10.**
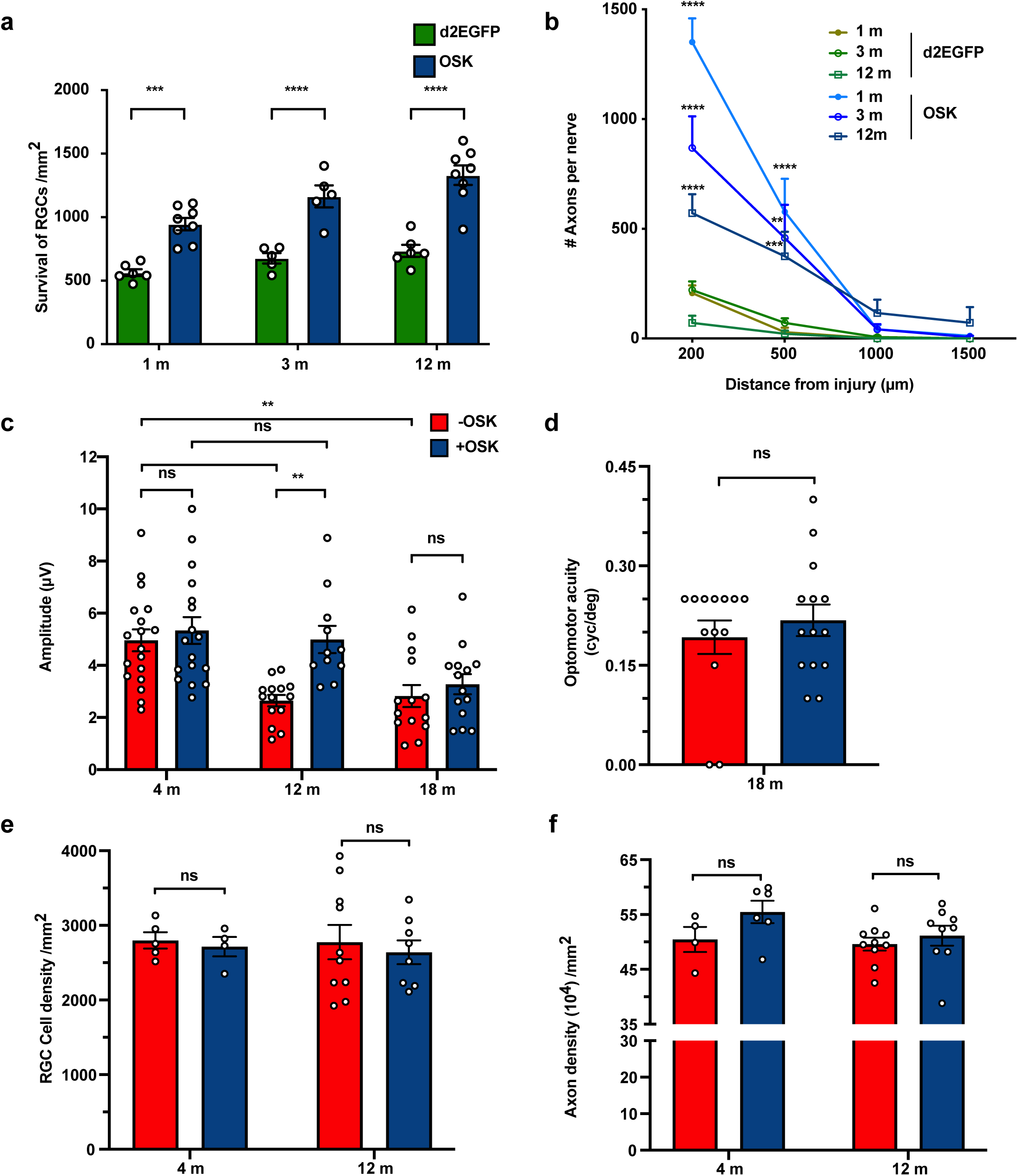
Effect of OSK in aged mice. **a**, Effect of OSK expression on RGC survival in young, adult, and aged mice after optic nerve crush injury. **b,** Axon regeneration after OSK expression, compared to the d2EGFP controls in young (1 m), adult (3 m), and aged (12 m) mice, 2 weeks post-injury. (The 12 m data in Fig.4b was reshown, as experiments were performed at the same time as the 1 m and 3 m old mice). **c,** Comparison of PERG measurement in different ages at one-month after -OSK or +OSK treatment. - OSK: AAV-rtTA+AAV-TRE-OSK; +OSK: AAV-tTA+AAV-TRE-OSK. **d**, Spatial frequency threshold in 18-month-old mice treated with -OSK or +OSK AAV for 4 weeks. **e,** Comparison of RGC cell density in 4- and 12 m-old-mice one month after -OSK or +OSK treatment. **f,** Comparison of axon density in 4 m- and 12 m-old mice, one month after -OSK or +OSK treatment. **p < 0.01, **** p < 0.0001, Two-way ANOVA with Bonferroni correction in **a**, **c**, **e**, **f.** One-way ANOVA with Bonferroni correction in **b.** Student’s t-test in **d**.

**Extended Data Fig. 11.**
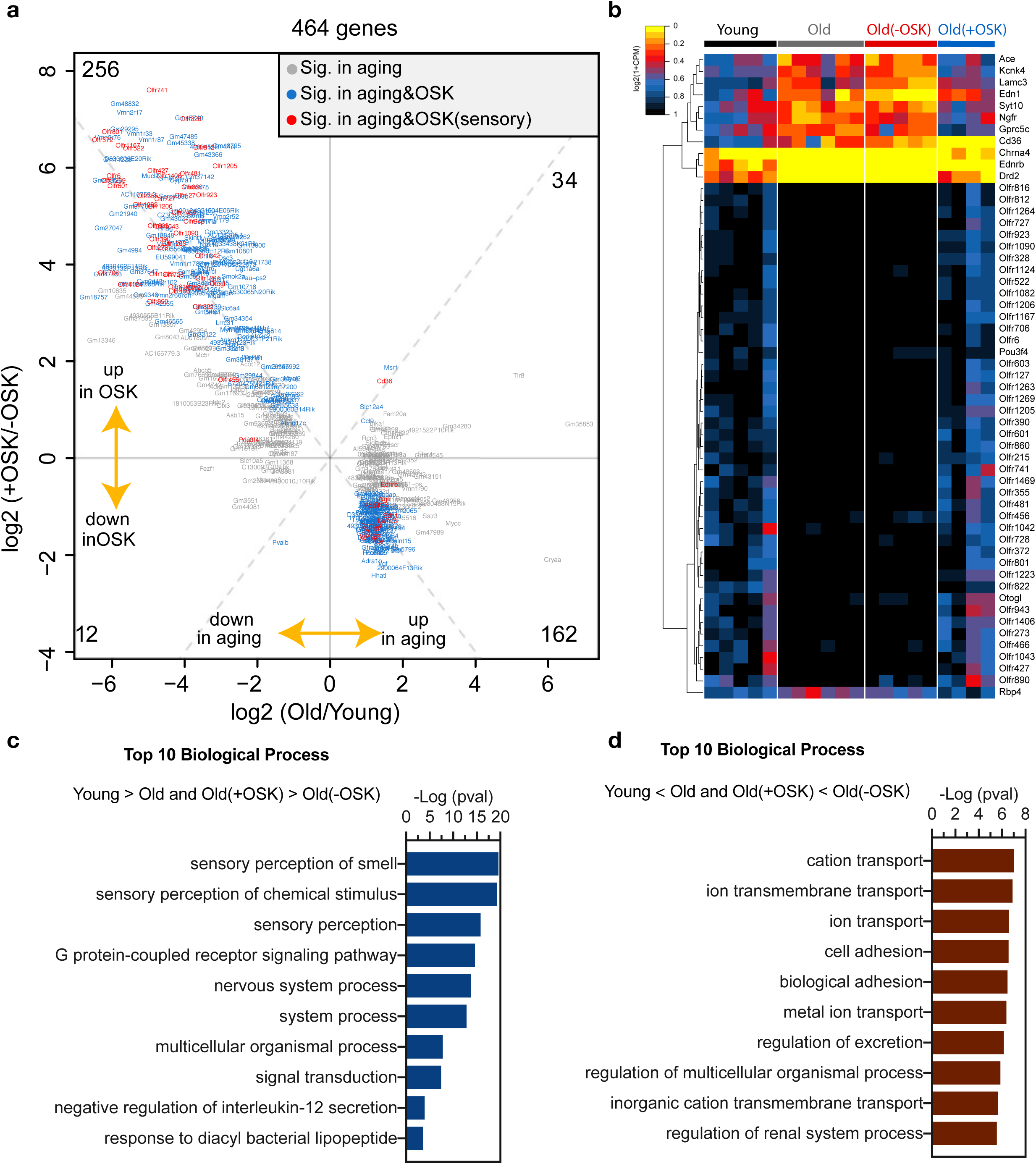
RNA-seq analysis of OSK-treated RGCs. **a**, Scatter plot of OSK-induced changes versus age-associated changes in mRNA levels. Dots represent differentially expressed genes in RGCs. Gene exclusion criteria: genes with low overall expression (log2(CPM)<2), genes that did not significantly change with age (absolute log2 fold-change <1) or genes altered by the virus (differentially expressed between intact old and old treated with TRE-OSK AAV). **b**, Hierarchical clustered heatmap showing RNA-Seq expression of sensory genes in FACS-sorted RGCs from young (5 m) or old mice (12 m), or old mice treated with either -OSK or +OSK AAV. **c**, Top 10 biological process that are lower in old compared to young and restored by OSK. **d**, Top 10 biological process that are higher in old compared to young and reduced by OSK.

## References

1. Sinclair, D. A., Mills, K. & Guarente, L. Accelerated aging and nucleolar fragmentation in yeast sgs1 mutants. Science 277, 1313–1316 (1997).

2. Oberdoerffer, P. & Sinclair, D. A. The role of nuclear architecture in genomic instability and ageing. Nat Rev Mol Cell Biol 8, 692–702, doi:10.1038/nrm2238 (2007).

3. Oberdoerffer, P. et al. SIRT1 redistribution on chromatin promotes genomic stability but alters gene expression during aging. Cell 135, 907–918, doi:10.1016/j.cell.2008.10.025 (2008).

4. Horvath, S. DNA methylation age of human tissues and cell types. Genome Biol 14, R115, doi:10.1186/gb-2013-14-10-r115 (2013).

5. Horvath, S. & Raj, K. DNA methylation-based biomarkers and the epigenetic clock theory of ageing. Nat Rev Genet 19, 371–384, doi:10.1038/s41576-018-0004-3 (2018).

6. Yun, M. H. Changes in Regenerative Capacity through Lifespan. Int J Mol Sci 16, 25392–25432, doi:10.3390/ijms161025392 (2015).

7. Goldberg, J. L., Klassen, M. P., Hua, Y. & Barres, B. A. Amacrine-signaled loss of intrinsic axon growth ability by retinal ganglion cells. Science 296, 1860–1864, doi:10.1126/science.1068428 (2002).

8. Waddington, C. H. The Strategy of the Genes; a Discussion of Some Aspects of Theoretical Biology London: George Allen & Unwin, Ltd. (1957).

9. Berger, S. L., Kouzarides, T., Shiekhattar, R. & Shilatifard, A. An operational definition of epigenetics. Genes Dev 23, 781–783, doi:10.1101/gad.1787609 (2009).

10. Mahmoudi, S., Xu, L. & Brunet, A. Turning back time with emerging rejuvenation strategies. Nat Cell Biol 21, 32–43, doi:10.1038/s41556-018-0206-0 (2019).

11. Takahashi, K. & Yamanaka, S. Induction of pluripotent stem cells from mouse embryonic and adult fibroblast cultures by defined factors. Cell 126, 663–676, doi:10.1016/j.cell.2006.07.024 (2006).

12. Petkovich, D. A. et al. Using DNA Methylation Profiling to Evaluate Biological Age and Longevity Interventions. Cell Metab 25, 954–960 e956, doi:10.1016/j.cmet.2017.03.016 (2017).

13. Ocampo, A. et al. In Vivo Amelioration of Age-Associated Hallmarks by Partial Reprogramming. Cell 167, 1719–1733 e1712, doi:10.1016/j.cell.2016.11.052 (2016).

14. Ohnishi, K. et al. Premature termination of reprogramming in vivo leads to cancer development through altered epigenetic regulation. Cell 156, 663–677, doi:10.1016/j.cell.2014.01.005 (2014).

15. Abad, M. et al. Reprogramming in vivo produces teratomas and iPS cells with totipotency features. Nature 502, 340–345, doi:10.1038/nature12586 (2013).

16. Gurdon, J. B. & Byrne, J. A. The first half-century of nuclear transplantation. Proc Natl Acad Sci U S A 100, 8048–8052, doi:10.1073/pnas.1337135100 (2003).

17. Shannon, C. E. A Mathematical Theory of Communication. The Bell System Technical Journal 27, 379–423 (1948).

18. Hofmann, J. W. et al. Reduced expression of MYC increases longevity and enhances healthspan. Cell 160, 477–488, doi:10.1016/j.cell.2014.12.016 (2015).

19. Smalley, E. First AAV gene therapy poised for landmark approval. Nat Biotechnol 35, 998–999, doi:10.1038/nbt1117-998 (2017).

20. Senis, E. et al. AAV vector-mediated in vivo reprogramming into pluripotency. Nat Commun 9, 2651, doi:10.1038/s41467-018-05059-x (2018).

21. Geoffroy, C. G., Hilton, B. J., Tetzlaff, W. & Zheng, B. Evidence for an Age-Dependent Decline in Axon Regeneration in the Adult Mammalian Central Nervous System. Cell Rep 15, 238–246, doi:10.1016/j.celrep.2016.03.028 (2016).

22. Moore, D. L. et al. KLF family members regulate intrinsic axon regeneration ability. Science 326, 298–301, doi:10.1126/science.1175737 (2009).

23. Qin, S., Zou, Y. & Zhang, C. L. Cross-talk between KLF4 and STAT3 regulates axon regeneration. Nat Commun 4, 2633, doi:10.1038/ncomms3633 (2013).

24. Park, K. K. et al. Promoting axon regeneration in the adult CNS by modulation of the PTEN/mTOR pathway. Science 322, 963–966, doi:10.1126/science.1161566 (2008).

25. Olova, N., Simpson, D. J., Marioni, R. E. & Chandra, T. Partial reprogramming induces a steady decline in epigenetic age before loss of somatic identity. Aging Cell 18, e12877, doi:10.1111/acel.12877 (2019).

26. Wang, M. & Lemos, B. Ribosomal DNA harbors an evolutionarily conserved clock of biological aging. Genome Res 29, 325–333, doi:10.1101/gr.241745.118 (2019).

27. Meer, M. V., Podolskiy, D. I., Tyshkovskiy, A. & Gladyshev, V. N. A whole lifespan mouse multi-tissue DNA methylation clock. Elife 7, doi:10.7554/eLife.40675 (2018).

28. Thompson, M. J. et al. A multi-tissue full lifespan epigenetic clock for mice. Aging (Albany NY) 10, 2832–2854, doi:10.18632/aging.101590 (2018).

29. Koh, K. P. et al. Tet1 and Tet2 regulate 5-hydroxymethylcytosine production and cell lineage specification in mouse embryonic stem cells. Cell Stem Cell 8, 200–213, doi:10.1016/j.stem.2011.01.008 (2011).

30. Gao, Y. et al. Replacement of Oct4 by Tet1 during iPSC induction reveals an important role of DNA methylation and hydroxymethylation in reprogramming. Cell Stem Cell 12, 453–469, doi:10.1016/j.stem.2013.02.005 (2013).

31. Guo, J. U., Su, Y., Zhong, C., Ming, G. L. & Song, H. Hydroxylation of 5-methylcytosine by TET1 promotes active DNA demethylation in the adult brain. Cell 145, 423–434, doi:10.1016/j.cell.2011.03.022 (2011).

32. Yu, H. et al. Tet3 regulates synaptic transmission and homeostatic plasticity via DNA oxidation and repair. Nat Neurosci 18, 836–843, doi:10.1038/nn.4008 (2015).

33. Weng, Y.-L. et al. An Intrinsic Epigenetic Barrier for Functional Axon Regeneration. Neuron 94, 337–346.e336, doi:10.1016/j.neuron.2017.03.034 (2017).

34. Horvath, S. et al. Epigenetic clock for skin and blood cells applied to Hutchinson Gilford Progeria Syndrome and ex vivo studies. Aging (Albany NY) 10, 1758–1775, doi:10.18632/aging.101508 (2018).

35. Heijl, A. et al. Reduction of Intraocular Pressure and Glaucoma Progression: Results From the Early Manifest Glaucoma Trial. JAMA Ophthalmology 120, 1268–1279, doi:10.1001/archopht.120.10.1268 (2002).

36. Krishnan, A. et al. Overexpression of Soluble Fas Ligand following Adeno-Associated Virus Gene Therapy Prevents Retinal Ganglion Cell Death in Chronic and Acute Murine Models of Glaucoma. J Immunol 197, 4626–4638, doi:10.4049/jimmunol.1601488 (2016).

37. Yao, K. et al. Restoration of vision after de novo genesis of rod photoreceptors in mammalian retinas. Nature 560, 484–488, doi:10.1038/s41586-018-0425-3 (2018).

38. McClellan, A. J. et al. Ocular surface disease and dacryoadenitis in aging C57BL/6 mice. Am J Pathol 184, 631–643, doi:10.1016/j.ajpath.2013.11.019 (2014).

39. Bar-Nur, O. et al. Small molecules facilitate rapid and synchronous iPSC generation. Nat Methods 11, 1170–1176, doi:10.1038/nmeth.3142 (2014).

40. Shipley, M. M., Mangold, C. A. & Szpara, M. L. Differentiation of the SH-SY5Y Human Neuroblastoma Cell Line. J Vis Exp, 53193, doi:10.3791/53193 (2016).

41. Triche, T. J., Jr., Weisenberger, D. J., Van Den Berg, D., Laird, P. W. & Siegmund, K. D. Low-level processing of Illumina Infinium DNA Methylation BeadArrays. Nucleic Acids Res 41, e90, doi:10.1093/nar/gkt090 (2013).

42. Fortin, J. P., Triche, T. J., Jr. & Hansen, K. D. Preprocessing, normalization and integration of the Illumina HumanMethylationEPIC array with minfi. Bioinformatics 33, 558–560, doi:10.1093/bioinformatics/btw691 (2017).

43. Dordea, A. C. et al. An open-source computational tool to automatically quantify immunolabeled retinal ganglion cells. Exp Eye Res 147, 50–56, doi:10.1016/j.exer.2016.04.012 (2016).

44. Gao, S. & Jakobs, T. C. Mice Homozygous for a Deletion in the Glaucoma Susceptibility Locus INK4 Show Increased Vulnerability of Retinal Ganglion Cells to Elevated Intraocular Pressure. Am J Pathol 186, 985–1005, doi:10.1016/j.ajpath.2015.11.026 (2016).

45. Sun, D., Qu, J. & Jakobs, T. C. Reversible reactivity by optic nerve astrocytes. Glia 61, 1218–1235, doi:10.1002/glia.22507 (2013).

46. Kim, D., Langmead, B. & Salzberg, S. L. HISAT: a fast spliced aligner with low memory requirements. Nat Methods 12, 357–360, doi:10.1038/nmeth.3317 (2015).

47. Liao, Y., Smyth, G. K. & Shi, W. featureCounts: an efficient general purpose program for assigning sequence reads to genomic features. Bioinformatics 30, 923–930, doi:10.1093/bioinformatics/btt656 (2014).

48. Robinson, M. D., McCarthy, D. J. & Smyth, G. K. edgeR: a Bioconductor package for differential expression analysis of digital gene expression data. Bioinformatics 26, 139–140, doi:10.1093/bioinformatics/btp616 (2010).

49. Carbon, S. et al. AmiGO: online access to ontology and annotation data. Bioinformatics 25, 288–289, doi:10.1093/bioinformatics/btn615 (2009).

50. Ashburner, M. et al. Gene ontology: tool for the unification of biology. The Gene Ontology Consortium. Nat Genet 25, 25–29, doi:10.1038/75556 (2000).

51. The Gene Ontology, C. The Gene Ontology Resource: 20 years and still GOing strong. Nucleic Acids Res 47, D330–D338, doi:10.1093/nar/gky1055 (2019).

